# Immune Checkpoint Molecules as Biomarkers of *Staphylococcus aureus* Bone Infection and Clinical Outcome

**DOI:** 10.1101/2024.12.30.630837

**Authors:** Motoo Saito, Katya A. McDonald, Alex K. Grier, Himanshu Meghwani, Javier Rangel-Moreno, Enrique Becerril-Villanueva, Armando Gamboa-Dominguez, Jennifer Bruno, Christopher A. Beck, Richard A. Proctor, Stephen L. Kates, Edward M. Schwarz, Gowrishankar Muthukrishnan

## Abstract

*Staphylococcus aureus* prosthetic joint infections (PJIs) are broadly considered incurable, and clinical diagnostics that guide conservative vs. aggressive surgical treatments don’t exist. Multi-omics studies in a humanized NSG-SGM3 BLT mouse model demonstrate human T cells: 1) are remarkably heterogenous in gene expression and numbers, and 2) exist as a mixed population of activated, progenitor-exhausted, and terminally-exhausted Th1/Th17 cells with increased expression of immune checkpoint proteins (LAG3, TIM-3). Importantly, these proteins are upregulated in the serum and the bone marrow of *S. aureus* PJI patients. A multiparametric nomogram combining high serum immune checkpoint protein levels with low proinflammatory cytokine levels (IFN-γ, IL-2, TNF-α, IL-17) revealed that TIM-3 was highly predictive of adverse disease outcomes (AUC=0.89). Hence, T cell impairment in the form of immune checkpoint expression and exhaustion could be a functional biomarker for *S. aureus* PJI disease outcome, and blockade of checkpoint proteins could potentially improve outcomes following surgery.

## Introduction

Chronic *Staphylococcus aureus* osteomyelitis, encompassing prosthetic joint infections (PJIs), is considered broadly incurable and has been the bane of orthopedic surgery^1^. Although the number of infections following elective orthopaedic surgery is low (1-5%), reinfection or relapse rates are very high (up to 30%) and cost up to $150,000 per patient^2^. Moreover, ∼13% of patients infected with *S. aureus* become septic and die from multiorgan failure, while others recover with relatively little intervention^3-5^. It is also known that patients can resolve acute infections and live full lives with asymptomatic *S. aureus* osteomyelitis^5-7^. Unfortunately, evidence-based clinical diagnostics to guide conservative vs. aggressive treatment of these patients do not exist, and no immunotherapies exist that can overcome the limitations of standard-of-care antibiotic treatment^8^. This led to an unprecedented 2018 International Consensus Meeting on musculoskeletal infections that concluded that “development of a functional definition for treatable “acute” vs. difficult-to-treat “chronic” osteomyelitis is the greatest research priority” in this field^1^. Thus, definitive empirical methods to discriminate acute vs. chronic-stage bone infections are a critical need for patient care, and to this end, a better understanding of the immune mechanisms that cause incurable *S. aureus* osteomyelitis is critical.

A conventional T cell response to acute *S. aureus* infection includes a burst in proliferation and differentiation upon activation, followed by the establishment of memory and contraction after pathogen clearance^9^. CD4 T cells, while responding to *S. aureus*, exhibit extensive plasticity, allowing the subsets to finely modulate one another while orchestrating local immunity through coordination of T-helper, cytotoxic, and immunosuppressive functions to curb hyperimmunity, cytokine storm, and thus prevent tissue damage^10-12^. T cell immunity studies in murine osteomyelitis models found that *S. aureus* skews the host-induced proinflammatory Th1 and Th17 responses during the early stages of infections, and then towards suppressive Treg responses in the late stages^13^, causing bacterial persistence in the bone. To study human T cell responses during *S. aureus* bone infections, we created a humanized NSG mouse model of chronic osteomyelitis, and demonstrated that the commencement of chronic implant-associated osteomyelitis occurs with large numbers of proliferating CD3^+^/Tbet^+^ adjacent to purulent abscesses in the bone marrow^14^. Interestingly, this coincided with increased infection and osteolysis, suggesting that human T cell infiltration in the bone does not aid with bacterial clearance at this stage. This suggests that these T cells exhibit dysfunction or impairment in the form of diminished effector function^15-18^ and cellular exhaustion^19-21^.

One of the known attributes of T cell dysfunction is exhaustion, which is well-characterized in chronic viral infections and cancer^19-22^. An exhausted T cell typically exhibits impaired effector function, reduced proliferative potential, increased expression of immune inhibitory receptors (e.g., LAG3^23,24^, TIM-3^25,26^, PD-1^27,28^, CTLA-4^29,30^), and altered cellular programming^22^. Our understanding of T cell exhaustion mechanisms during *S. aureus* infections is limited. However, it is known that *S. aureus* superantigens (SAgs) can trigger antigen-independent oligoclonal T cell activation and proliferation^31-33^, leading to secretion of high amounts of proinflammatory cytokines^15-18^, followed by a profound state of T cell exhaustion characterized by lack of proliferation, cytokine production, and apoptosis^34-37^. Nonetheless, whether T cell exhaustion occurs in a human chronic osteomyelitis setting remains to be investigated.

In the current study, we utilize humanized NSG-SGM3 mice engrafted with the human fetal liver and thymus^38^, human patient samples, and multi-omics to comprehensively examine CD4 T cell exhaustion in the bone niche during chronic *S. aureus* osteomyelitis. Importantly, we investigated whether immune checkpoint expression and exhaustion could be utilized as biomarkers of adverse disease outcomes in patients with *S. aureus* osteomyelitis.

## Results

### Humanized NSG-SGM3 BLT mice exhibit exacerbated *S. aureus* implant-associated osteomyelitis

Our previous studies revealed increased susceptibility of humanization of NSG mice to *S. aureus* osteomyelitis, which included exacerbated suppuration and sepsis^14^. However, the NSG mouse model has inherent limitations, including: 1) lack of functional thymic environment that supports the human T cell development and 2) limited myeloid lineage development with diminished macrophage function^39,40^. Therefore, we generated an improved humanized mouse model of osteomyelitis with NSG-SGM3 mice expressing human KITLG, GM-CSF, and IL-3 to allow for enhanced human myeloid lineage development^41,42^. These animals were subjected to sublethal radiation-induced myeloablation and transplanted with donor-matched human CD34+ HSC fetal liver cells and thymic tissues under the mice kidney capsule to generate human immune cells and improve T cell development (**Figure 1A**). At 12 weeks post-engraftment, humanized NSG-SGM3 BLT mice were assessed for the extent of human chimerism as described previously^14^.

**Figure 1.**
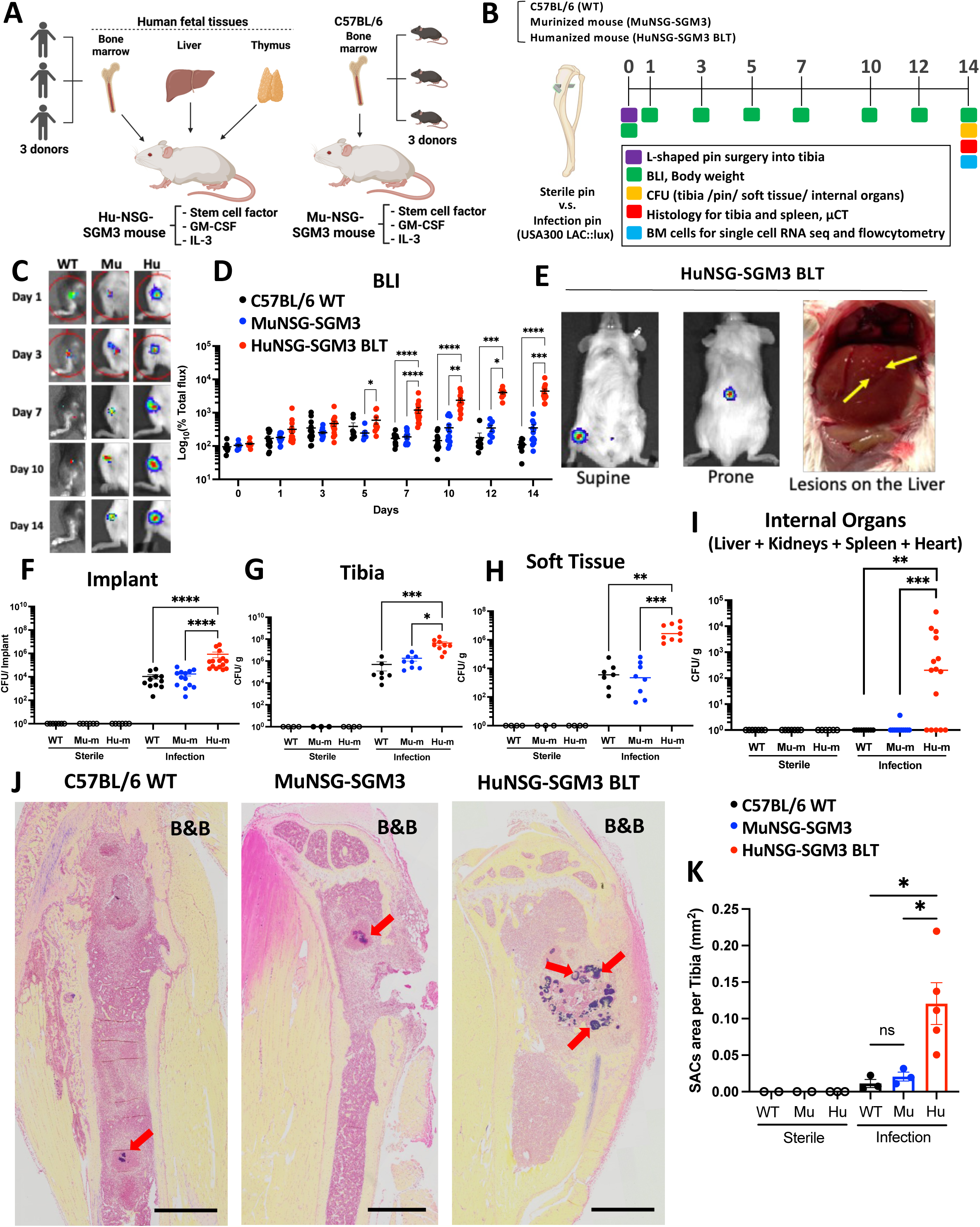
Humanized NSG-SGM3 BLT mice have exacerbated susceptibility to *S. aureus* osteomyelitis compared to Murinized NSG-SGM3 and C57BL/6 WT mice. (**A**) Humanized NSG-SGM3 BLT mice were generated by engrafting with CD34^+^ human hematopoietic cells, autologous human fetal liver, and thymus from **three different human donors**. Murinized NSG-SGM3 BLT mice were generated with CD34^+^ murine hematopoietic cells derived from three different C57BL/6 WT mice. (**B**) Schematic illustration of the experimental design of in vivo experiments. 20-week-old humanized HuNSG-SGM3 BLT mice, murinized NSG-SGM3 and C57BL6 (WT) mice (n=25) were subjected to transtibial implant-associated osteomyelitis using bioluminescent MRSA (USA300 LAC::*lux*). (**C**) Longitudinal BLI images of representative mice with (**D**) statistical analysis of the groups demonstrate increased in vivo *S. aureus* growth in humanized NSG-SGM3 BLT mice. (**E**) In vivo BLI images of a representative NSG-SGM3 BLT mouse with local and disseminated MRSA infections, as evidenced by the focal BLI signal in the tibia and abdominal cavity from supine and prone views, respectively. Autopsy photograph confirmed *S. aureus* abscesses (yellow arrows) in the liver. (**F-I**) On day14 post-operation, implants, tibiae, surrounding soft tissues, and internal organs (heart, liver, kidneys, and spleen) were harvested for CFU assays and the data are presented with the mean for each group (n= 25, and differences between groups were assessed by ANOVA, *p<0.05, **p<0.01, ***p<0.001, ***p<0.0001). (**J**) Representative 10x images of Brown & Brenn (B&B) stained histology of infected tibia from each group are shown, highlighting the SACs (red arrows). (**K**) VisioPharm histomorphometry was performed to quantify the SAC area per tibia, and the value for each tibia is presented with the mean +/-SD (n≥4, ANOVA, *p<0.05).

Subsequently, we examined MRSA (USA300 LAC::*lux*) implant-associated osteomyelitis in these animals using our established protocols^14,43-47^. We hypothesized that infection would be more severe in humanized NSG-SGM3 BLT mice (Hu-m) compared to murinized NSG-SGM3 (Mu-m) (engrafted with C57/BL6 bone marrow CD34+ HSCs) and C57BL6 (WT) control mice (**Figure 1A-B**). The results demonstrated that infected BLT mice experienced increased in vivo *S. aureus* growth vs. controls as measured by bioluminescence **(Figure 1C-D).** Additionally, BLT mice had increased infection severity in the bone niche and sepsis **(Figure 1E-I)**. This included internal organ dissemination **(Figures 1E and 1I)**, an over 45-fold increase in CFUs on the pin/implant (WT= 1.09 x 10^4^, Mu-m= 1.75 x 10^4^, Hu-m= 8.67 x 10^5^), an over 20-fold increase in CFUs/g in the bone (WT= 6.51 x 10^5^, Mu-m= 2.3 x 10^6^, Hu-m= 5.29 x 10^7^), and an over 450-fold increase in CFUs/g in the soft tissue (WT= 1.17 x 10^4^, Mu-m= 1.32 x 10^4^, Hu-m=6.33 x 10^6^) **(Figure 1F-H).** Lastly, histopathology analyses of Brown & Brenn-stained tibiae sections revealed that the BLT mice exhibited increased staphylococcal abscess communities (SAC) compared to control animals **(Figure 1J-K**, *p<0.05). These results confirmed increased susceptibility of humanized BLT mice to *S. aureus* osteomyelitis.

### The human T cell landscape in the bone niche revealed remarkable heterogeneity during *S. aureus* osteomyelitis

We previously observed large numbers of proliferating human CD3^+^/Tbet+ cells in the bone niche of humanized CD34+ NSG mice^14^. To comprehensively elucidate the human T cell landscape in the bone microenvironment during osteomyelitis, we performed single-cell RNAseq analysis of tibial bone marrow cells isolated from MRSA-infected NSG-SGM3 BLT mice at 2 weeks post-infection (**Figure 2A**). Specifically, bone marrow cells were isolated from MRSA-infected and sterile implant control humanized BLT mice tibias and sorted into human CD45^+^CD3^+^ T cells and CD45^+^CD19^+^ B cells, mixed at 1:1 proportion and subjected to scRNAseq analyses (**Figure 2A**). Approximately 30,000 cells were sequenced, and unsupervised clustering analyses were performed using R Studio Seurat packages (v4.0.3)^48-51^ **(Figure S1)**. A total of 39 clusters were revealed, which, upon re-clustering, were segregated into 24 T cell clusters and 16 B cell clusters (**Figure 2B-C**). T cell clustering data were normalized to the total number of T cells to account for human donor-to-donor variability. Subsequent UMAP^52^ clustering of the identified human T cell clusters revealed remarkable heterogeneity in gene expression (**Figure 2D-E**) and T cell population numbers (**Figure 2E**) between sterile and infection surgery groups. Notably, the number of Th1/Th17 cells (red arrow, clusters 8 and 20) was prominently increased in the infected animals compared to sterile implant controls, suggesting that human Th1/Th17 responses predominate due to *S. aureus* in the bone niche at later stages of infection (2-weeks post-surgery), indicative of persistent osteomyelitis.

**Figure 2.**
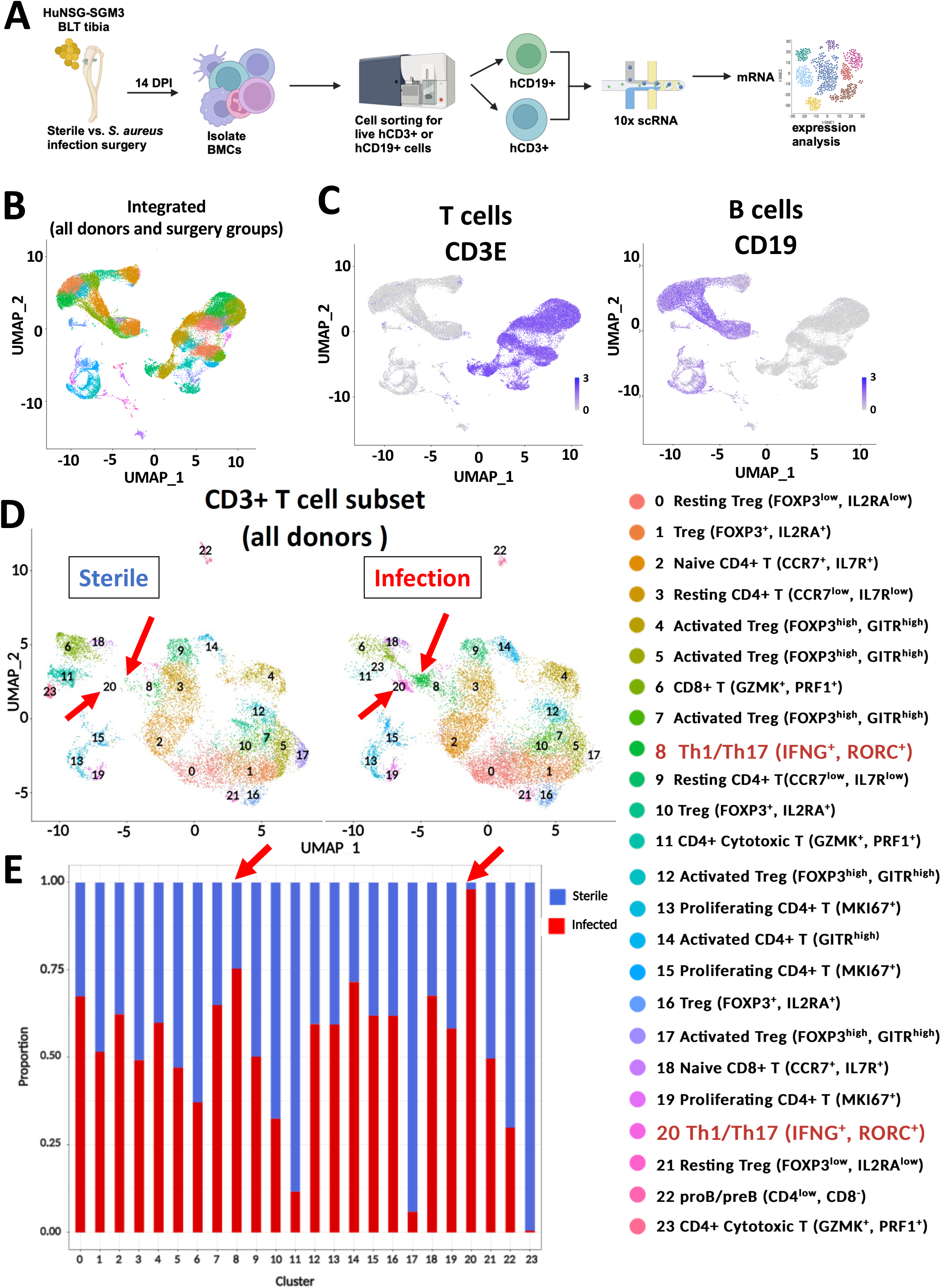
Single-cell RNAseq reveals remarkable human T cell heterogeneity at the infection site in humanized BLT mice with *S. aureus* osteomyelitis. (**A**) Schematic illustration showing the experimental overview of sc-RNAseq of humanized NSG-SGM3 BLT mice engrafted with three different human donor tissues. Bone marrow (BM) cells were collected from tibiae of humanized NSG-SGM3 BLT mice 14 days after transtibial implants surgery with or without USA300 LAC::*lux*, and the human CD45^+^CD19^+^ B cells and CD45^+^CD3^+^ T cells were isolated by FACS for scRNAseq. (**B**) UMAP of the unsupervised cluster analysis of ∼30,000 BM cells with (**C**) Feature plots of the CD3^+^ T cells and CD19^+^ B cells. (**D**) UMAP and DEG clustering analyses of hCD45^+^/CD3^+^ T cells identified 24 T cell clusters with (**E**) bar graphs displaying the proportion of cell counts in each cluster between sterile implant and infected implant groups. Note the marked increase of Th1/Th17 cells (red arrows, Cluster 8,20) in the infected tibiae compared to unifected tibiae.

### Immune checkpoint proteins are elevated in the CD4+ T cells in *S. aureus*-infected humanized BLT mice tibia

Human Th1/Th17 cells (clusters 8 & 20) were sub-clustered to reveal 7 clusters, and differential expression of gene (DEG) analyses were performed to probe for immune activation and suppression genes (**Figure 3A**). Interestingly, several of these clusters showed significantly increased expression of immune checkpoint proteins LAG-3, and TIM-3 (HAVCR2) in the infected animals (**Figure 3B**). Importantly, transcriptional factor TCF1 (TCF7), known to be associated with “progenitor-exhausted” cells in CD8 T cells, were up-regulated in some of the Th1/Th17 clusters. TOX and TOX2, which are associated with functional terminal exhaustion, were up-regulated in other T cell clusters (**Figure 3C**). Of note, we observed higher expression of CXCL13 in the infected compared to controls, and the CXCL13/CXCR5 axis has recently been implicated in driving CD8 T cells to the progenitor-exhausted phenotype^53^. We also observed diminished proliferative (MKi67) and cytokine-producing capacities in these cells (**Figure 3C**).

**Figure 3:**
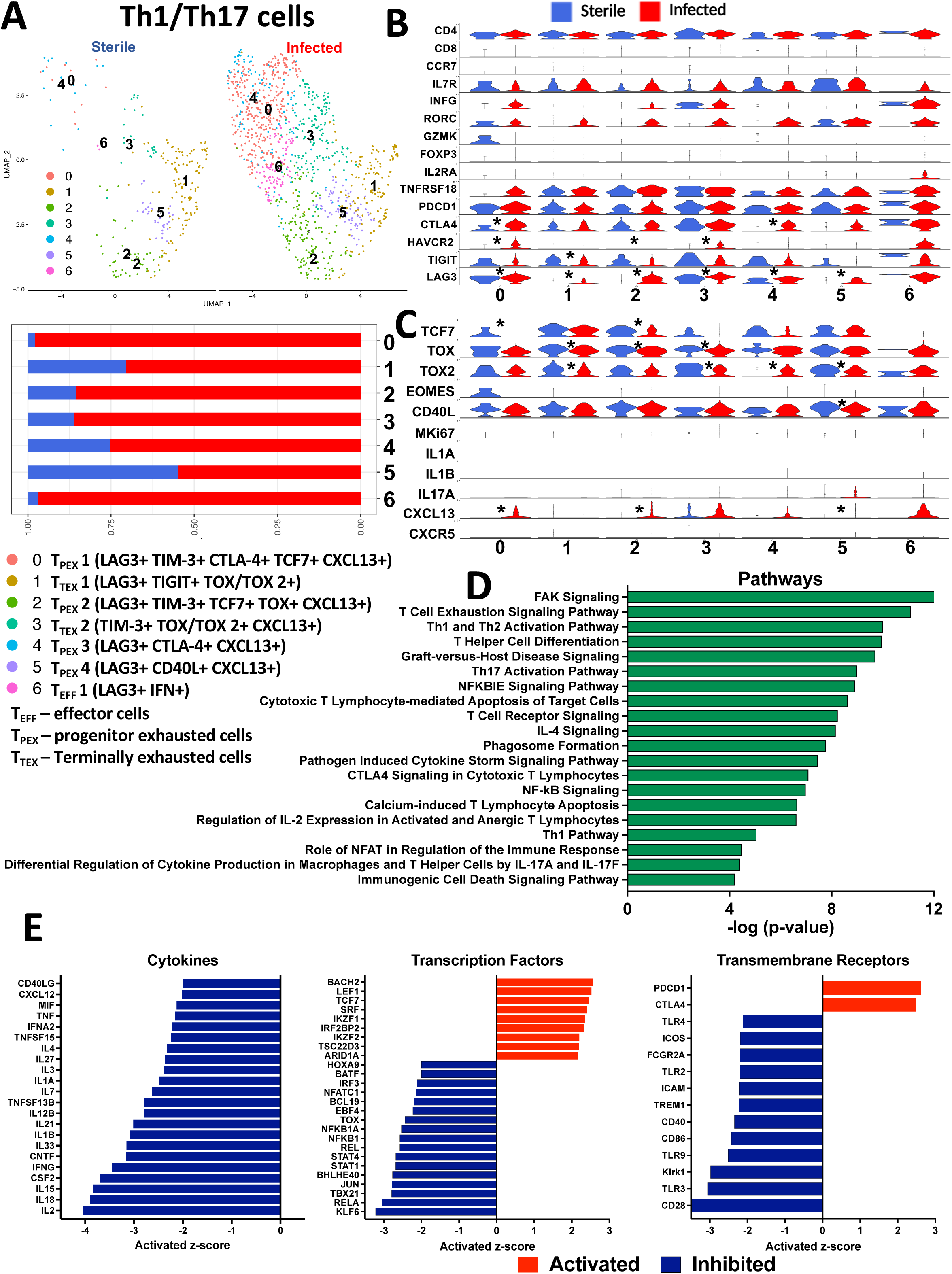
Immune checkpoint gene expression is elevated in CD4+ Th1/Th17 cells from *S. aureus*-infected humanized BLT tibiae. **(A)** The scRNAseq data of the Th1/Th17 cells (clusters 8 and 20) identified in Figure 2 were subjected to UMAP and differential gene expression analyses (DEG) revealed 7 sub-clusters, and the relative proportions of these sub-clusters in uninfected (blue) and infected (red) tibiae are illustrated by the bar graph. (**B**) Violin plot analyses demonstrated that these cells were of the Th1/Th17 phenotype. Several Th1/Th17 clusters showed significantly increased expression of immune checkpoint molecules LAG-3, TIM-3 (HAVCR2), and, to a lesser extent, CTLA-4 and other immunosuppressive genes like TIGIT. (**C**) DEG analyses of transcriptional factors (TCF7, TOX1-2, EOMES, NR4A1), cytokines & chemokines, and chemokine receptor (IL-1, IL-17, CXCL13, CXCR5) associated with functional T cell exhaustion, chronic antigenic stimulation (CD40L) and proliferation (MKi67). Note that the lower expression of TCF7, MKi67, IL-1, and IL-17 genes and higher expression of CXCL13 and TOX 2 indicate transcriptional reprogramming of these cells to a terminally functionally exhausted state (*p<0.05). The Th1/Th17 subclusters were annotated based on the gene expression signatures into activated, progenitor-exhausted, and terminally-exhausted cells. The DEGs between the experimental groups within the Th1/Th17 cells were subjected to Ingenuity Pathway Analysis (IPA) to identify the (**D**) top significantly enriched canonical pathways and (**E**) predicted upstream regulators (cytokines, transcriptional factors and transmembrane receptors). Red indicates activation, while blue indicates suppression.

Next, the DEGs between the experimental groups within the Th1/Th17 cluster of cells were subjected to Ingenuity Pathway Analysis (IPA) to identify the top significantly enriched canonical pathways and predicted upstream regulators **(Figure 3D-E)**. Notably, IPA confirmed that the T cell exhaustion signaling pathway was one of the top 3 significantly enriched pathways **(Figure 3D)**. Additionally, predicted upstream-activated proteins included transmembrane receptors such as CTLA-4, PDCD1, and transcriptional factor TCF7 **(Figure 3E)**, which are associated with exhaustion. Furthermore, inhibited upstream proteins included multiple cytokines (IL2, IFNG, TNF, and IL7), TLRs (TLR4, TLR2, TLR3, and TLR9), and transcription factors (NFKB1, STAT1, STAT4, and IRF3), suggesting diminished effector functions **(Figure 3E)**. Collectively, these results suggest that human CD4+ Th1/Th17 cells likely undergo functional exhaustion at the chronic stage of osteomyelitis.

### Evidence of functional cellular exhaustion in human T cells at the bone infection site in BLT mice

To confirm scRNAseq findings at the protein level, we next performed immunohistochemistry (IHC) on the tibiae of infected and uninfected humanized BLT mice. IHC confirmed the presence of LAG3^+^, TIM-3^+^, and PD-1^+^ T cells clustering next to SACs (**Figure 4A**) in the MRSA-infected animals. Importantly, spectral flow cytometric analyses revealed that the frequency of human CD3^+^CD4^+^ T cells expressing TIM-3, LAG-3 & PD-1 in tibiae from MRSA-infected BLT mice were significantly higher compared to controls (**Figure 4B, Supplemental Figure S2A**). Next, we examined the responses of these CD3^+^CD4^+^ T cells following in vitro stimulation with PMA and ionomycin via spectral flow cytometry **(Figure 5A).** We observed that unstimulated CD4^+^TIM-3^+^ and CD4^+^LAG3^+^ cells had a significantly lower frequency of proliferating Ki67^+^ cells compared to CD4^+^TIM-3^-^ and CD4^+^LAG-3^-^ cells in the infected samples, suggesting that these cells are functionally exhausted **(Figure 5B)**. We also evaluated cytokine production by CD4^+^CD69^+^ T cells post-stimulation **(Supplemental Figure S2B)** and looked for differences between LAG-3^+^ and LAG-3^-^ cells and TIM-3^+^ and TIM-3^-^ cells within this population. In general, we observed that MRSA infection induced more cytokine-producing CD4^+^CD69^+^ cells (**Supplemental Figure S3**). The results showed that TIM-3^+^ cells made significantly less TNFα and IL-17A, and a trending decrease in IFN-γ and IL-2 compared to TIM-3-cells **(Figure 5C, supplemental Figure S4)**. Similarly, LAG-3^+^ cells made significantly less IL-2 and IL-17a and a trending decrease in IFN-γ compared to LAG-3^-^ cells **(Figure 5C).** Of note, examination of splenocytes revealed similar trends of decreased proliferative capacity and diminished effector functions, suggesting systemic effects of infection (**Supplemental Figure S5**). These results demonstrate the impaired functional capacity of LAG3^+^ and TIM-3^+^ cells in our model, providing further evidence of T cell exhaustion during chronic osteomyelitis.

**Figure 4.**
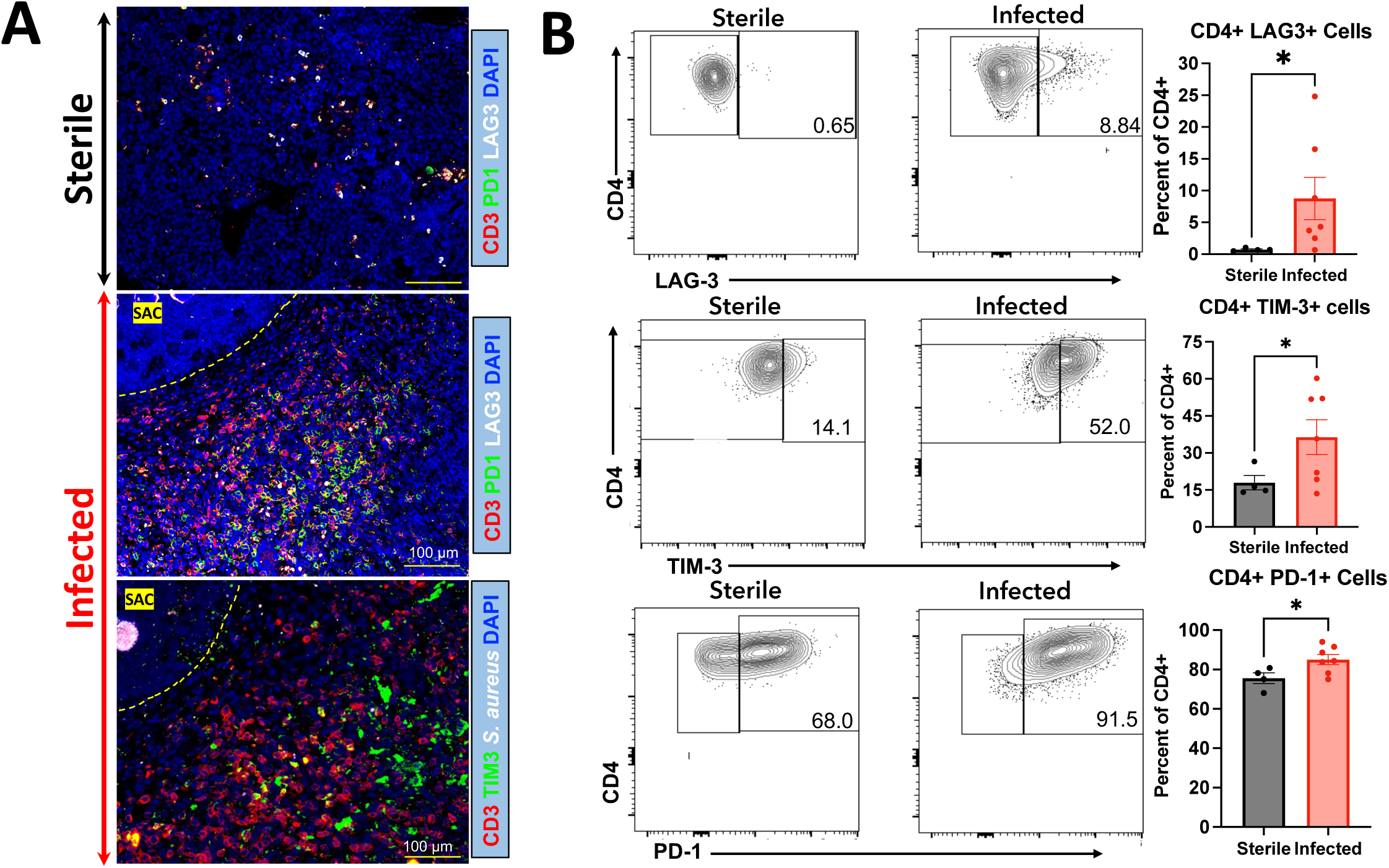
CD4+ T cells expressing immune checkpoint proteins are increased in *S. aureus*-infected humanized BLT tibiae. **(A)** Immunofluorescent histochemistry analyses of tibia sections from uninfected and MRSA-infected humanized BLT mice 14 days post-op were performed with labeled antibodies against CD3, LAG-3, TIM-3, and PD-1 with DAPI counter stain, and representative images are shown at 4x. Note the increased numbers of T cells near the SAC (dashed yellow line) in the infected tibiae. **(B)** A multichromatic spectral flow cytometry analyses were performed on tibial bone marrow cells from uninfected and MRSA-infected BLT mice. Live human CD45+/CD3+/T cells and their subpopulations (CD4+, CD8+, Tregs) were analysed for immune checkpoint expression (LAG3, TIM-3, and PD-1) and proliferation (Ki67), and representative histograms are shown. Note the frequency of human CD3^+^/CD4^+^ T cells expressing TIM-3, LAG3 & PD-1) in the cells from MRSA-infected bone marrow (n=4-8 mice, *p<0.05, t-test).

**Figure 5:**
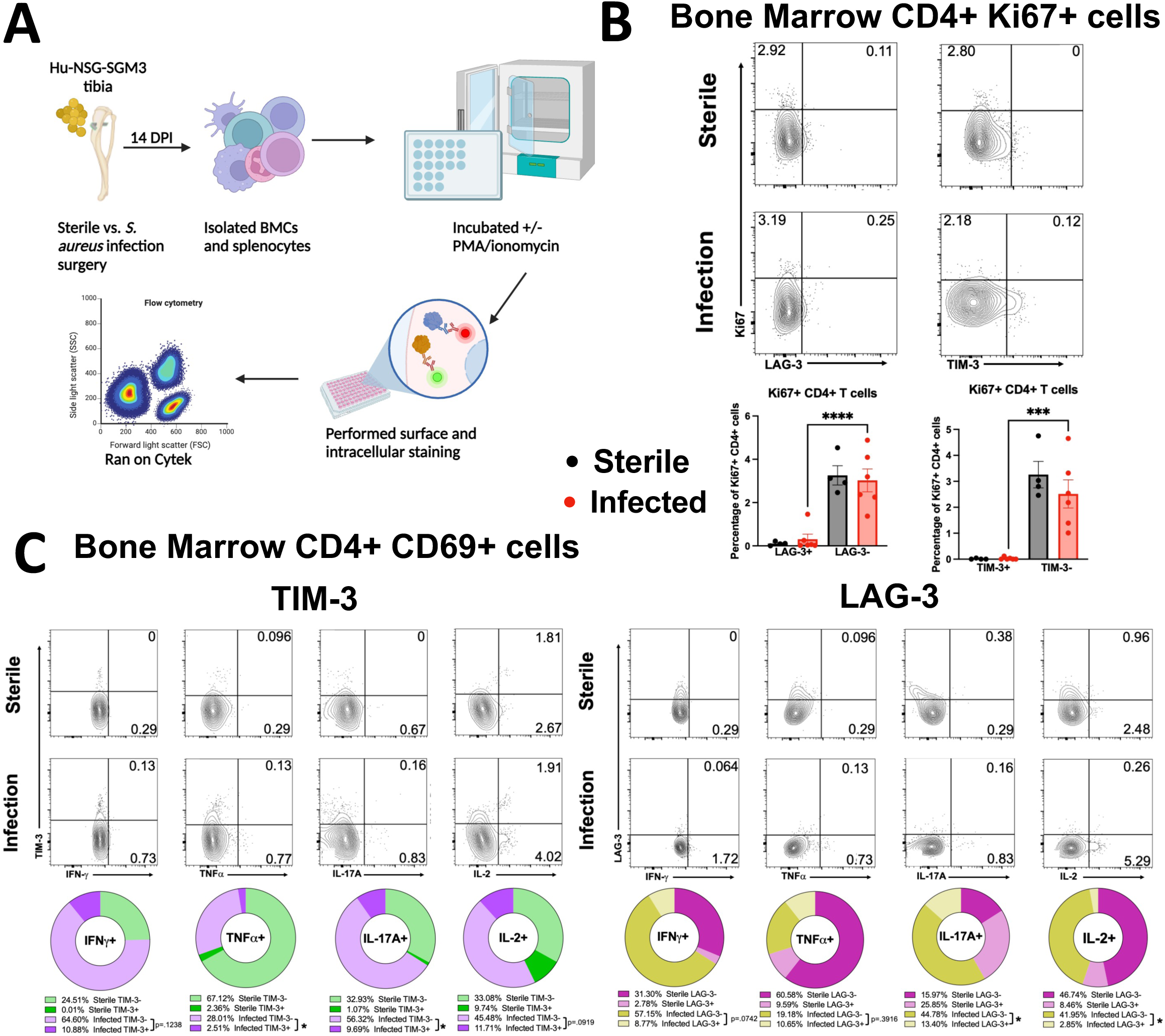
Bone marrow CD4+ T cells from MRSA-infected tibiae expressing TIM-3 and LAG3 checkpoint proteins exhibit diminished proliferative capacity and altered cytokine production. **(A)** Schematic illustration of the experimental design of ex-vivo experiments. 20-week-old humanized NSG-SGM3 BLT mice were subjected to aseptic or septic transtibial implant-surgery for 14 days, then their splenocytes and bone marrow cells were isolated, stimulated, stained with antibodies, and analyzed by flow cytometry. **(B-C)** Multichromatic spectral flow cytometry was performed on uninfected and MRSA-infected tibial bone marrow cells from BLT mice **(B)** on unstimulated cells **(C)** post-stimulation with PMA/ionomycin. **(B)** Live human CD45+/CD3+/CD4+ T cells expressing checkpoint molecules TIM-3 and LAG-3 were probed for their proliferative capacity using the cell surface marker Ki67. Note that CD4+TIM-3+ and CD4+LAG-3+ cells have lower amounts of proliferating Ki67+ cells in the bone marrow of infected BLT mice, suggesting functional exhaustion and dysfunction. **(C)** Live human CD45+/CD3+/CD4+/CD69+ T cells expressing checkpoint molecules TIM-3 and LAG-3 were probed for functional capacity using the cytokines IFN-!, TNFɑ, IL-17A, and IL-2 (n=4-9 mice, *p<0.05, ANOVA).

### Immune checkpoint protein expression in the human bones of *S. aureus* PJI patients

Next, we examined whether our observations from the humanized mouse model were predictive of human immunity in patients with *S. aureus* PJI. To test this, we performed histology on bone sections from *S. aureus*-infected patients. H&E staining revealed considerable immune infiltration at the site of infection **(Figure 6A)**. IHC showed co-expression of immune checkpoint proteins in CD3^+^ T cells and CD66b^+^ neutrophils **(Figure 6B-D)**. Specifically, PD-1^+^CD3^+^ **(Figure 6B)**, TIM-3^+^CD3^+^ **(Figure 6C)**, and LAG-3^+^CD3^+^ **(Figure 6C)** cells were observed. We also observed TIM-3^+^ neutrophils **(Figure 6D)**. These results indicate that human bones infected with *S. aureus* provide an environment that is supportive of T cell exhaustion during PJI.

**Figure 6.**
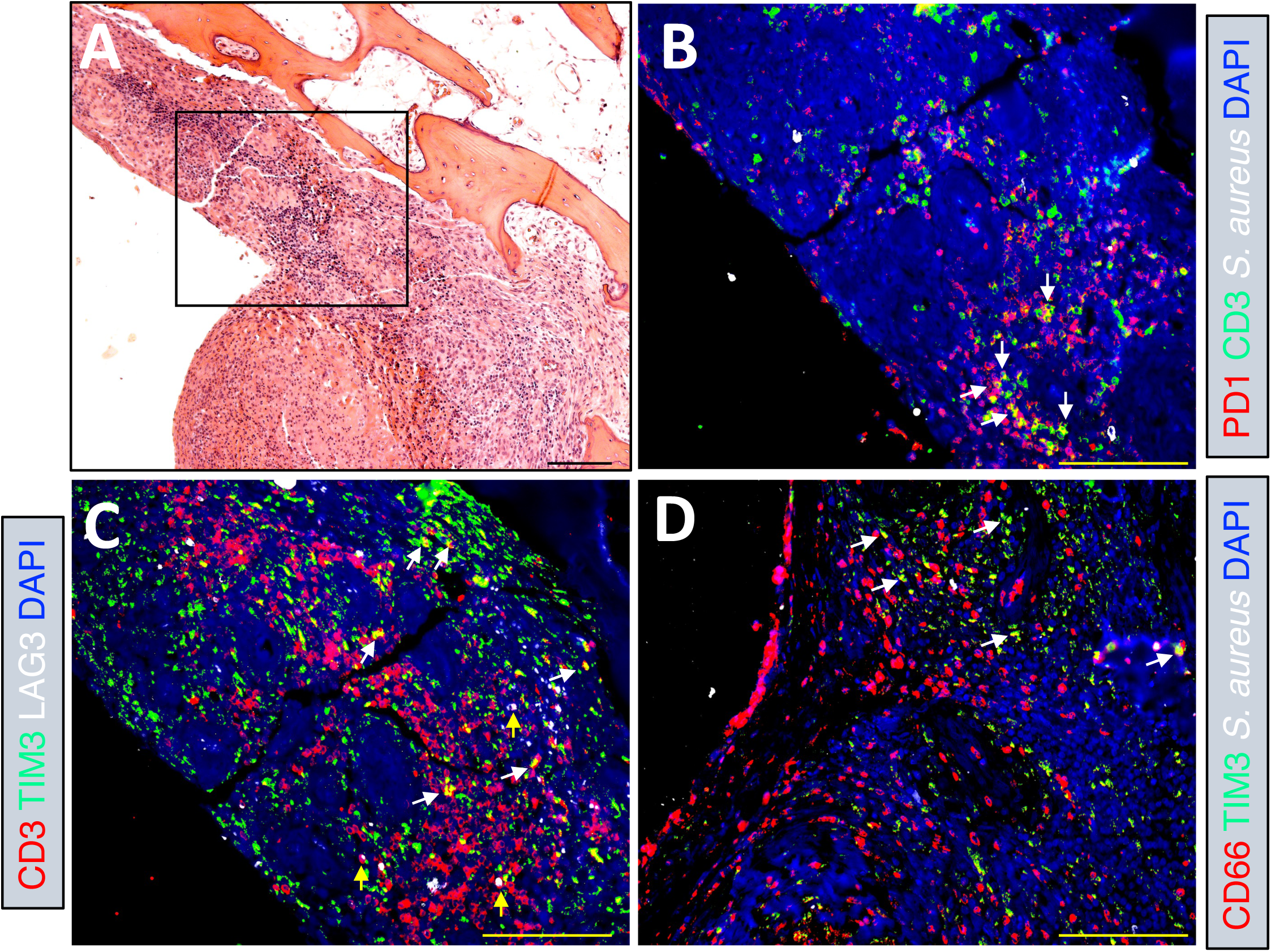
T cells expressing immune checkpoint proteins accumulate in *S. aureus* infected bone tissue from PJI patients. Bone tissues surgically removed from PJI patient with *S. aureus* osteomyelitis were processed for histology and immunohistochemistry. **(A)** Representative 100x image (bar =100 μm) of a H&E-stained section is shown to illustrate the inflammatory cells within the region of interest (box). (**B-D)** Parallel histology sections containing the region of interest were immunostained with labelled antibodies against CD3, PD1, *S. aureus,* TIM-3 (green), LAG-3, and CD66b, counter stained with DAPI, and representative fluorescent microscopy images are shown at 200x (bar = 100 μm). (**B**) Note CD3^+^/PD1^+^ T cells detected in areas of *S. aureus* infection (white arrows). **(C)** Note CD3^+^/TIM-3^+^ (white arrows) and CD3^+^/LAG-3^+^ (yellow arrows) T cells at the site of *S. aureus* infection. **(D)** Note TIM-3^+^/CD66b+ neutrophils at the site of infection (white arrows).

### Serum immune checkpoint proteins are prognostic of adverse outcomes in *S. aureus* osteomyelitis patients

Next, we assessed these immune checkpoint proteins in the serum of orthopaedic patients with culture-confirmed *S. aureus* osteomyelitis and individuals undergoing total hip/knee arthroplasties with no infections (**Figure 7A**, **supplemental Table 1**). The serum samples collected prior to surgery were examined. Serum LAG3 levels were significantly upregulated in *S. aureus* patients compared to uninfected individuals. Moderate trending elevations in TIM-3 and CTLA-4 were observed in the infected patients (**Figure 7B**). Multivariate logistic regression analyses with risk characterized by odds ratios (OR per 10-fold increase in protein levels) revealed that TIM-3 levels were significantly associated with adverse outcomes (OR = 485.1, 95% CI 2.49 -94511.09, p = 0.02) such as arthrodesis, reinfection, amputation, and septic death. A multiparametric nomogram (TEX) combining TIM-3 with LAG3, PD-1, and CTLA-4 was highly predictive of adverse outcomes (AUC=0.89) in osteomyelitis patients (**Figure 7E-F**). No correlation was observed with the anecdotal clinical classification of acute vs. chronic disease (**Figure 7C-D**). Our results suggest these proteins could be leveraged as prognostic biomarkers for *S. aureus* osteomyelitis treatment outcomes.

**Figure 7.**
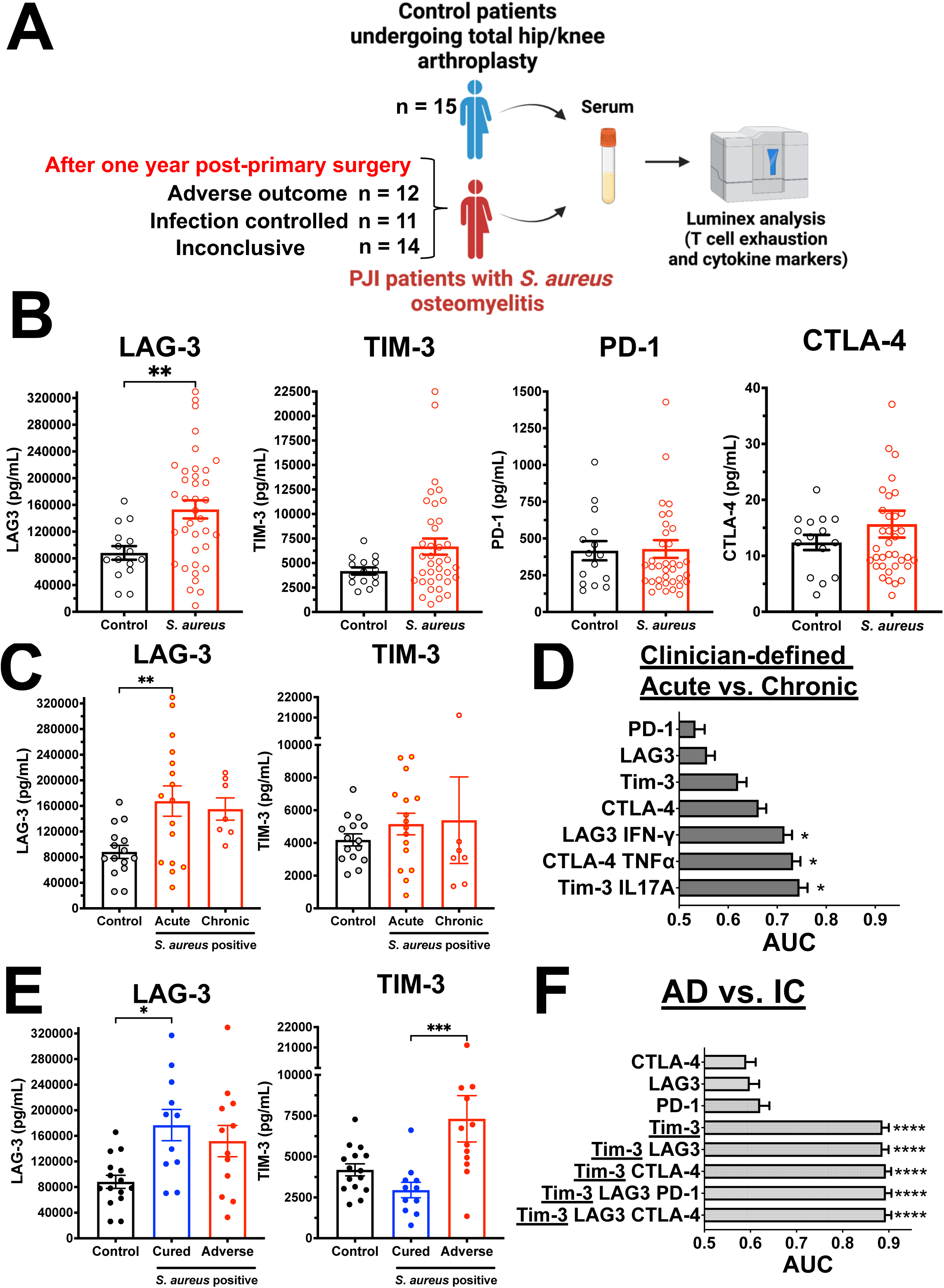
TIM-3 protein level in serum is highly prognostic of adverse outcomes in patients with *S. aureus* osteomyelitis. **(A)** Serum samples were collected from healthy arthritis patients undergoing total hip/knee arthroplasty (n=15), and orthopaedic patients undergoing surgery for culture-confirmed *S. aureus* osteomyelitis whose clinical outcome at 1-year was adverse (n=12), infection controlled (n=11), or inconclusive (14). (**B**) Immune checkpoint proteins LAG-3, TIM-3, CTLA-4, PD-1 and cytokines (IFN-γ, IL-2, TNFα, IL-17A, IL-17F) were assessed by multiplex Luminex assay, and the data are presented for each patient with the mean +/-SEM for each group. The individual protein levels were utilized to perform receiver operating characteristic (ROC) curve analysis either singly or in combination to generate the area under the curve (AUC) for **(C-D)** differentiating acute vs. chronic *S. aureus* infections and (**E-F**) prognostic prediction of outcome. Interestingly, no correlation was observed between levels of immune checkpoint proteins and clinical time-based, anecdotal classification of acute vs. chronic classification. On the other hand, immune checkpoint proteins, especially TIM-3, were highly predictive of adverse in these patients (*p<0.05, **p<0.01, ****p<0.00001).

## Discussion

Clinical diagnostics that guide aggressive vs. conservative treatments for serious bone infections do not exist. To this end, we studied a humanized mouse model of osteomyelitis and human patient samples to assess if T cell exhaustion could be the much-needed evidence-based prognostic for disease outcome.

Identifying a biomarker predictive of adverse outcomes during bone and joint infection has been a priority in musculoskeletal research. It has led to considerable efforts evaluating the effectiveness of antibodies for *S. aureus*-specific antigens^54^, cytokines^55^, and host-derived proteins^56^. Here, we demonstrate that immune checkpoint proteins in patient serum were markedly increased in adverse outcomes compared to uninfected and cured patients. Our human serum data illustrate the potential for checkpoint proteins as a biomarker of adverse outcomes in hip and knee arthroplasty and may provide empiric data upon which to base clinical decisions.

MRSA infection of NSG-SGM3 BLT mice in the bone resulted in an exacerbated infection phenotype characterized by increased bacterial load in the bone, MRSA dissemination to distant internal organs, and purulent abscess formation compared to non-humanized mice. Our group and others have extensively observed this increased severity of *S. aureus* in osteomyelitis, pneumonia, bacteremia, soft-tissue, and deep-tissue abscess infections^14,57-61^. These studies highlight the potential involvement of immunotoxins and virulence proteins that exhibit high tropism to human leukocytes^62,63^. For instance, SAgs exhibit a 100 to 1000-fold decreased mitogenic activity in murine and rat-derived T cells compared to human T cells^15-17,64,65^.

The importance of T cells in chronic *S. aureus* infection control has been investigated in mice and humans^11^. Specifically, studies have observed that *S. aureus* can induce an immunostimulatory Th1/Th17 response, which can transition to immunosuppressive Tregs over time if the host fails to clear the infection^13^. These Tregs have been implicated in broader T cell suppression, along with myeloid-derived suppressor cells (MDSCs)^66^. Additionally, in overall bone health, the balance of Th17s and Tregs is important in determining osteoclastogenesis^67^. Expectedly, we observed an increase in Th1/Th17 cells and moderate increases in Treg populations at 2 weeks post-infection in our humanized mouse model.

A more detailed examination of Th1/Th17 cells revealed that these cells may exist in a mixed population of “activated,” “progenitor exhausted,” and “terminally exhausted” cells. T cell exhaustion occurs in multiple tissues and organs during chronic viral infection, inflammation, or cancer^22^. However, evidence during bacterial infections is less clear. Studies have shown the occurrence of CD4 and CD8 exhaustion in the periphery during tuberculosis infection, but the local site of infection has not been evaluated^68^. Notably, a recent study examining human PJI tissue T cells and PMNs revealed enrichment of T cell exhaustion signaling pathways and immune suppression, such as PD-1/PD-L1 pathways^69^, consistent with our observations in humanized mice and human patient tissue samples. In addition to local bone niche, we observed splenic CD4 T cell exhaustion, suggesting that *S. aureus* can induce systemic T cell dysfunction. Our findings corroborated a recent study in chronic osteomyelitis patients with *S. aureus* and non-*S. aureus* infections^70^. The authors observed systemically increased Tregs and Tfh populations in the peripheral blood of infected patients compared to uninfected controls. Curiously, they observed moderate increases in PD-1 and TIM-3 expression in B cells, dendritic cells, and monocytes but not in T cells^70^. Nonetheless, such studies highlight cellular exhaustion beyond T cells during osteomyelitis, and that exhaustion is likely caused by pathogens beyond *S. aureus* in a chronic infection setting.

An important consideration in developing and maintaining T cell function is the impact of other cell types, such as macrophages, in the bone marrow local infection site. *S. aureus* has been known to manipulate T cell responses by inducing hypoxia and supporting MDSC development, which leads to T cell suppression ^11,71^. Indeed, MDSCs are key contributors to *S. aureus* orthopedic biofilm infections^72,73^, and our IHC revealed some non-T cell PD-1+ cells in the infected humanized mice, which could be MDSCs. Moreover, T cell activities in the bone niche due to infection can further diminish oxygen, exacerbate hypoxia^74,75^, and ultimately promote *S. aureus* biofilm formation, typical of chronic *S. aureus* disease. It is also known that exhausted CD8 T cells promote MDSC formation^76^, and it remains to be investigated if exhausted CD4 T cells also do this.

*S. aureus*-infected macrophages in the bone marrow aid in apoptotic cell clearance, ultimately leading to hypoxia^77,78^. This efferocytosis of apoptotic cells by macrophages inhibits their antigen presentation and has been shown to contribute to skewing naïve T cells into Treg fate^79^. Our scRNAseq and IPA analyses revealed up-regulated exhaustion, apoptotic pathways in Th1/Th17 cells, and a substantial population of Tregs. Interestingly, exhaustion-indicative checkpoint expression has been observed on the CD4 Tregs in the tumors of glioblastoma patients^80^. Nonetheless, further work is needed to understand the dynamics between these cells during *S. aureus* infection.

We observed CD4 T cell exhaustion 2 weeks post-infection, which is considered the initiation of chronic infection with a set inoculation dosage. As exhaustion is driven by antigenic stimulation^81^, the initial bacterial load may impact the development of this dysfunction. This has been shown using transgenic mouse models but not with a pathogen^82^. More work is warranted into how pathogen loads influence the T cell response in PJI. This is especially relevant in bacterial infections, where a single causative agent can lead to a spectrum of disease states^83^.

The observation of T cell exhaustion in the local infection niche naturally leads to the idea of using immune checkpoint blockade (ICB) therapies. They are routinely used in the clinic to treat specific cancer malignancies, including melanoma, non-small cell lung cancer, and esophageal squamous cell carcinoma^84^. The use of immunotherapy in *S. aureus* infections has been proposed previously. A group used anti-PD-L1 as an adjuvant with gentamycin during *S. aureus* osteomyelitis and found an improved histological score and decreased cortical bone loss at fourteen days post-infection^85^. However, such therapies should identify the ideal balance between reenergizing T cells to clear the infection and preventing excess tissue damage due to inflammation. Indeed, there is evidence from human PJI patients that prolonged pro-inflammatory immune responses can hinder bone healing^86^. Future immunotherapies against this disease cannot adopt a one-size-fits-all approach. Importantly, treatment responses should also be tailored to the tissue microenvironment, as a recent study demonstrated that the local tissue niche has profound effects on immune and metabolic responses to *S. aureus*^69^.

The current study has a few limitations. First, we only utilized a single time point in our murine infection studies, and it will be important to examine the temporal changes of CD4 T cell responses and exhaustion over time. Understanding the kinetics of the T cell response is crucial to determining the potential of immune checkpoint proteins as biomarkers or the use of ICB therapies. Additionally, we used naïve humanized mice in our infection studies, which meant a lack of immunological memory. As most humans have been exposed to *S. aureus*^87^, the impact of memory T cells on the development of exhaustion during osteomyelitis and how it influences disease pathogenesis is unknown*. S. aureus*-specific memory CD4 T cells have been observed in human skin, but how they may respond to pathogenic challenge by *S. aureus*, or if they exist in other tissues, has not yet been investigated^88^. Additionally, as effector memory cells can become exhausted post-reactivation, further investigation into the contributions of naïve and memory cells to the observed T cell exhaustion is warranted^89^. Finally, our clinical pilot study examining serum immune checkpoint proteins and outcomes was limited in sample size. A larger, more comprehensive prospective study is required to establish that these proteins are definitive biomarkers of adverse outcomes due to *S. aureus* osteomyelitis.

## Materials and Methods

### Ethics Statement

The University Committee on Animal Resources and the Institutional Animal Care and Use Committee have reviewed and approved all animal husbandry and experimentation. All mice were maintained in an Accreditation of Laboratory Animal Care (AAALAC) International-approved facility.

### Human Patient Samples

*<u>For Luminex Immunoassays:</u>* Recruited patients were either enrolled in an international biospecimen registry (AO Trauma Clinical Priority Program (CPP) Bone Infection Registry^90^) or participated in IRB-approved clinical studies at Virginia Commonwealth University^91^. Patient information was collected in a REDCap database managed by AO Trauma and VCU data management administrators. Laboratory investigators had access to the serum of patients and their deidentified clinical data, which was provided on request by the data management teams.

*<u>For Immunofluorescence Analyses</u> in_Human Bones with Microbial Abscesses:* Human bones with histological or microbiological evidence of bacterial infection were provided by Drs. Enrique Becerril-Villanueva and Armando Gamboa-Dominguez (n = 3). Specimen collection was conducted with written and signed consent from their family members in accordance with the Declaration of Helsinki and after approval from the Ethical Committee of the national Institute of Medical Sciences and Nutrition “Salvador Zubiran”. Analysis of deidentified tissue samples was performed according to protocols approved by the University of Rochester Review Board. Patients received a diagnosis of a bone infection from experienced microbiologists and pathologists.

Among the three patients identified, Patient 1 (female, 51 years, type 2 diabetic) was diagnosed with an *S. aureus* abscess in the metatarsus, had a visible wound without exposed bone, and was not undergoing antibiotic treatment. Patient 2 (male, 28 years, type 1 diabetic) and Patient 3 (male, 45 years, type 1 diabetic), undergoing insulin treatment for diabetes management, were diagnosed with *S. capitis/S. haemolyticus* and *E. coli* osteomyelitis and had visible wounds with exposed metatarsal bone. Note that patients 2 and 3 were undergoing antibiotic therapy with Erythromycin + Dicillin and Dicillin, respectively, at the time of the biopsy.

Patient biopsies were collected from a patient hospitalized for amputation of the first metatarsal of the left foot; the sample was collected in paraformaldehyde 10% for a subsequent decalcification process, and the sample was embedded in paraffin and then sectioned at a thickness of 5 μm. H&E staining showed acute ulcerated inflammation with the presence of osteomyelitis and mononuclear cells associated with the presence of large positive bacteria. Patient 1 data is presented in Figure 6.

### Murine model of implant-associated osteomyelitis

*<u>Mouse strains/humanization:</u>* Female C57BL/6J mice (stock 000664) and NSG-SGM3 (NOD.Cg-Prkdc^scid^ Il2rg^tm1Wjl^ Tg(CMV-IL3,CSF2,KITLG)1Eav/MloySzJ stock 013062) mice were purchased from the Jackson Laboratories (Bar Harbor, ME, USA), housed five per cage in two-way housing on a 12-h light/dark cycle, and fed a maintenance diet and water ad libitum. Humanized and murinized NSG-SGM3 mice were provided by the Humanized Mouse Core (HMC) Facility, CCTI, CUMC, Columbia University. Humanized mice were generated by engrafting NSG-SGM3 mice with CD34+ human hematopoietic stem cells from fetal liver and thymic tissues according to previously described protocols^59,92-94^. Briefly, NSG-SGM3 mice (4 week) were subjected to total body irradiation (1 Gy) and injected intravenously with lineage-negative human CD34^+^ hematopoietic stem cells (2 x 10^5^ cells/mice) isolated from fetal liver tissue. Thymic tissue was also implanted under the kidney capsule. At 12 weeks post engraftment, mice were subjected to submandibular bleeding to isolate peripheral lymphocytes, and human immune cell reconstitution was assessed by flow cytometry as described previously^14^. In this study, tissue samples from a total of seven human male and female human donors were utilized for generating humanized mice, and we obtained 61.2% ± 21.8% human CD45+ cell engraftment (<u>human T– 36.5%±11.7% human B– 35.5% ± 22.1%</u>). Murinized mice were generated by engrafting NSG-SGM3 mice with lineage-negative C57BL/6J CD34+ bone marrow cells.

*<u>MRSA Infection Studies:</u>* Transtibial implant-associated osteomyelitis with MRSA was performed on skeletally mature 20–24-week-old humanized NSG-SGM3 mice, and age-matched C57BL/J6 and murinized NSG-SGM3 mice utilizing our well-validated protocols described previously^14,47,95^. A bioluminescent strain of USA300 LAC (USA300 LAC::lux) was used in the infection studies^43,47,96,97^. Briefly, mice were anesthetized with isoflurane in a Plexiglass box (ca. 7% in O2, flow rate 0.6-1 L/min), maintained with isoflurane through a face mask (ca. 2-3% in O2, flow rate 0.6-1 L/min). Peri- and postoperative analgesia consisted of buprenorphine extended release, which was given subcutaneously prior to surgery (25mg/L). Before surgery, a flat stainless-steel surgical wire (cross-section, 0.2 mm by 0.5 mm) 4 mm long (MicroDyne Technologies, Plainville, CT, USA) bent at 1mm to form an L-shape was steam sterilized and used for sterile implant controls, or inoculated with clinical *S. aureus* USA300 LAC::lux strain grown overnight. After anesthesia induction, the right leg was clipped, and the skin was aseptically prepared with chlorhexidine scrub (Hibiscrub, 4% Chlorhexidine Digluconate) and 70% ethanol. The implant localization was identified (2 to 3 mm under the tibial plateau in the proximal tibia) using the proximal patella as an anatomical landmark and the jaws of the Mayo-Hegar needle driver as the measure. A hole was pre-drilled in the proximal tibia using a percutaneous approach from the medial to lateral cortex using a 26-gauge needle. Subsequently, *S. aureus* infected pin (5.0 x 10^5^ colony forming units (CFU)/mL) was surgically implanted in the pre-drilled hole from the medial to the lateral cortex. Osteotomy and implant position were confirmed radiographically in the lateral plane immediately after surgery. At 14 days post-infection, mice were euthanized, and the infected leg containing the transtibial implant was excised out for either CFU quantitation, histology, flow cytometry, or single-cell RNA sequencing. Additionally, internal organs, including the liver, spleen, kidneys, and heart, were harvested for CFU enumeration. Murine infection studies were performed four independent times, and the results shown are pooled data from these experiments.

*<u>Euthanasia:</u>* The tibia, tibial implant, and soft tissue abscesses surrounding the tibia were removed, weighed, and placed in 1mL or 2mL of room-temperature sterile PBS. The implant was sonicated for 15 minutes to dislodge attached bacteria, and organ tissues were homogenized (Omni TH, tissue homogenizer TH-02/TH21649, Kennesaw, GA, USA) in 2mL of PBS. Implant sonicate fluid and tissue homogenates were serially diluted, plated on blood agar (BA) plates, and incubated overnight at 37°C. To confirm *S. aureus* on the plates, random colonies from each plate/organ/tissue were picked, and StaphLatex agglutination test (Thermo Fisher Scientific, Waltham, MA, USA) was performed. Bacterial colonies were enumerated, and the generated CFU data were presented as CFUs per gram of tissue.

### Histopathology

The tibia was dissected from mice post-euthanasia and fixed for 72 hours in 4% neutral buffered formalin. Each mouse tibia was then rinsed with ddH2O and decalcified in 14% EDTA tetrasodium solution for 14 days at room temperature. Following decalcification, samples were paraffin-embedded, cut into 5 μm transverse sections, and mounted on glass slides for histological staining. Slides were deparaffinized and stained with Brown and Brenn (Gram) staining as described previously^14^. Digital images of the stained slides were created using VS120 Virtual Slide Microscope (Olympus, Waltham, MA, USA). Numbers SACs were manually enumerated and averaged across two or more histologic sections at least 50 μm apart from 6-7 mice in each experimental group. Quantitative analysis of SAC area within the tibias of C57BL/6J WT, murinized NSG-SGM3, and humanized NSG-SGM3 animals was performed on Brown and Brenn (Gram) stained slides using Visiopharm (v.2019.07; Hoersholm, Denmark) colorimetric histomorphometry utilizing a custom Analysis Protocol Package (APP). Manual regions-of-interest (ROIs) were drawn around the tibia and SACs within the tibia on each image prior to batch processing for automated quantification of SAC area normalized to tibial area between the groups.

### Multicolor Immunofluorescence

*<u>Primary antibodies:</u>* The following antibodies were utilized for immunostaining: goat anti-CD3ε (clone M-20, sc-1127, RRID:AB_631128, Santa Cruz Biotechnology), mouse anti-PD-1 (10377-MM23, RRID:AB_2936309, Sino Biologicals), Rabbit anti-LAG3 (clone BLR027F, NBP2-76402, RRID:AB_3403543, Novus Biologicals), Mouse anti-TIM3/HAVCR2 (clone TIM3/4031, V8754-20UG, NSJ Bioreagents), Rabbit anti-*S. aureus* (PA1-7246, RRID:AB_561546, Thermo Fisher Scientific), and Mouse anti-CD66b (G10F5, NBP2-80664, RRID:AB_3096017, Novus Biologicals).

*<u>Secondary antibodies:</u>* The following antibodies were used at 1:200 dilution for the detection and visualization of primary antibodies: Alexa Fluor 568-conjugated donkey anti-goat IgG (A-11057, RRID: AB_2534104, Thermo Fisher Scientific) to detect CD3-epsilon, Alexa Flour 488-conjugated donkey anti-rabbit IgG (711-546-152, RRID:AB_2340619, Jackson ImmunoResearch Laboratories) at a 1:200 dilution for detecting LAG-3 and *S. aureus*, Cy3-goat anti-mouse IgM (115-165-020, RRID:AB_2338683, Jackson ImmunoResearch Laboratories) to detect CD66b, FITC-donkey anti-mouse IgG (715-095-150, RRID:AB_2340792, Jackson ImmunoResearch Laboratories) to visualize PD1, Alexa Fluor 647 donkey anti-mouse Ig G (715-606-150, RRID: AB_2340865, Jackson ImmunoResearch Laboratories) to detect TIM-3.

*<u>Staining Procedure:</u>* The 5 μm formalin-fixed paraffin sections were incubated at 60°C overnight for deparaffinization. Tissue sections were quickly transferred to xylene and gradually hydrated by transferring slides to absolute alcohol, 96% alcohol, 70% alcohol, and then water. Slides were immersed in an antigen retrieval solution, boiled for 30 minutes, and cooled down for 10 minutes at room temperature (RT). Slides were rinsed several times in water and transferred to PBS. Non-specific binding was blocked with 5% normal donkey serum in PBS containing 0.1% Tween 20, 0.1% Triton-X-100 for 30 minutes, at RT in a humid chamber. Primary antibodies were added to slides and incubated in a humid chamber at RT, ON. Slides were quickly washed in PBS, and fluorescently labeled secondary antibodies were incubated for 2 hours at RT overnight in a humid chamber. Finally, slides were rinsed for 1 hour in PBS and mounted with Vectashield antifade mounting media with DAPI (H-1200, Vector Laboratories, Burlingame, CA, USA). Pictures were taken with a Zeiss Axioplan 2 microscope and recorded with a Hamamatsu camera.

### Cell Culture

Single-cell suspensions were generated from spleens and tibias. Following euthanasia, spleens were harvested and collected in 2mL PBS, then transferred through a 70µM filter. Spleens were resuspended in 2mL ACK lysis buffer (ThermoFisher, catalog: A1049201), and then filtered through a 40µm strainer. Bone marrow cells were isolated by flushing the bone with 1mL of PBS. For phenotyping analysis, cells were frozen and stored in liquid nitrogen, and then thawed. For functional analysis, immediately post-isolation, 1 x 10^6^ cells were stimulated (2uL/1mL, eBioscience Cell Stimulation Cocktail, ThermoFisher, catalog: 00-4970-93 in R10A2 media (RPMI (ThermoFisher, catalog: 11875093) + 10% FBS (ThermoFisher, catalog: 26130079) + antimycotic/antibiotic (ThermoFisher, catalog:15240062)) or unstimulated for 10 hours at 37°C 5% CO^2^, then left in 4°C overnight.

### Flow Cytometry

Two panels were used to interrogate changes in the immune response. Immunophenotyping of spleen and bone marrow from humanized NSG-SGM3 mice was performed. Briefly, for our immunophenotyping panel, single-cell suspension of splenocytes and bone marrow cells were thawed, and for our functional panel, cells were taken after stimulation. For both panels, 10^6^ cells/mouse were initially stained with fixable viability dye eFluor™ 780 (eBioscience™, Thermo Fisher Scientific catalog: 65-0865-18) for 30 minutes at 4° C to exclude dead cells from the analysis. Following washing and blocking with 5% normal mouse serum (ThermoFisher, catalog: 10410), surface antibody cocktails were added for 80 minutes at 4° C (Supplemental Tables X). After additional washing, cells were fixed/permeabilized with the BD Cytofix/Cytoperm Fixation/Permeabilization Kit and blocked again with normal mouse serum (BD Biosciences, catalog 554714). Intracellular antibody cocktails (Supplemental Tables X) were added for 80 minutes at 4° C. After staining, the cells were fixed with 2% formaldehyde before running on a Cytek Aurora five-laser spectral flow cytometer (Cytek Biosciences). Flow data were analyzed using FlowJo version 10.6 (Tree Star Inc. Ashland, OR). For both panels, single-color compensation controls for these antibodies were created using UltraComp eBeads Plus Compensation beads (Thermo Fisher Scientific, catalog 01-3333-43). All antibodies were purchased from BioLegend, BD Biosciences (San Jose, CA, USA), or Thermo Fisher Scientific.

### Single-cell RNA sequencing

Tibias from NSG-SGM3 BLT mice that underwent surgery with or without bioluminescent MRSA-contaminated transtibial implant were harvested on day 14 post-infection. Bone marrow cells were isolated in PBS. The cell suspension was stained with viability dye 7-AAD (BD Biosciences Cat# 559925, RRID:AB_2869266), anti-CD45 (Biolegend, catalog: 268530, RRID: AB_2715890), anti-CD19 (Biolegend, catalog: 353006, RRID: AB_2564128), and anti-CD3 (BD Biosciences, catalog: 555332, RRID: AB_395739). Isolated BM cells were sorted into human CD45+CD19+ B cells and CD45+CD3+ T cells on FACS Aria (BD Biosciences). Equal proportions of the B and T cells were subjected to scRNAseq analyses. A total # of events were collected for each sample and processed for single-cell RNA sequencing by the Genomics Research Center at the University of Rochester Medical Center. The cells were then sequenced using Illumina’s NovaSeq6000.

The datasets were analyzed using the Seurat^51^ package version 3 in R with unsupervised SNN clustering and the Louvain method. After initially clustering all cells at a resolution of 2.4 (determined by evaluating clustree plots^98^), T cells were extracted based on SingerR^99^ annotations and the absence of CD19. T cell clustering was performed at resolution = 1.0 (determined by evaluating clustree plots), using 30 principal components based on the top 2,000 variable genes. For analysis, low-quality cells were removed if the cell exhibited 1) <1000 genes, 2) >7000 genes, 3) >50,000 mapped reads, or 4) >5% mitochondrial reads. Integration of samples was done using the CCA method with SCTransform^100^ normalization, regressing out the effect of percent mitochondrial reads. We used the Human Primary Cell Atlas to identify gene expression patterns used to manually annotate each T cell cluster’s cell type. Sub-clustering of Th1/Th17 cells was performed at a resolution of 0.2, using 30 principal components based on the expression of exhaustion-associated genes. Differential expression analysis was performed using Seurat’s FindMarkers function with default parameters.

### Luminex Immunoassay

Serum concentrations of immune checkpoint proteins (TIM-3, LAG-3, PD-1, PD-L1, PD-L2, CTLA-4) and cytokines (IFN-γ, IL-2, TNF-α, IL-17A, and IL-17F) were determined in individuals undergoing total hip/knee arthroplasty and orthopaedic patients with culture-confirmed *S. aureus* osteomyelitis using a Luminex-based Milliplex xMAP Multiplex Assay (MilliporeSigma) according to the manufacturer’s instructions.

### Statistics

Unpaired student’s t-test was used to compare the flow cytometry data statistically. Two-way ANOVA with Sidak’s post-hoc tests were performed to compare body weight change over time. One-way ANOVA analyses with Tukey’s post-hoc tests were utilized to compare the osteolysis area, number of SACs, SAC area, log-transformed CFUs, and the number of immune cells revealed by immunostaining. The individual protein levels from patient serum samples in the clinical pilot stody were utilized to perform receiver operating characteristic (ROC) curve analysis either singly or in combination to generate the area under the curve (AUC) for differentiating acute vs. chronic *S. aureus* infections and prognostic prediction of outcome. All data and statistical analyses were conducted using GraphPad Prism (version 9.0), SAS version 9.4, and R Studio Seurat packages, and p < 0.05 was considered significant.

### Reporting summary and Data Availability

All scRNAseq datasets generated are available via the Gene Expression Omnibus (GEO) under the accession code GSE269658. All other data will be made available upon reasonable request. Further information on research design is available in the Nature Portfolio Reporting Summary linked to this article.

## Acknowledgments

This work was made possible by a University Research Award from the University of Rochester, NIH NIAMS P50 AR07200 Pilot Award with additional funding from the AOTrauma Clinical Priority Program, NIH (P30 AR069655 and P50 AR07200), and AAI Careers in Immunology fellowship program. The authors thank the Electron Microscope Shared Resource Laboratory, Genomics Research Center, and the Histology, Biochemistry, and Molecular Imaging Core in the Center for Musculoskeletal Research at the University of Rochester Medical Center.

## Figure Legends

**<u>Supplementary Figure S1.</u> Quality Control, Dimension Reduction, and Clustering Overview.** (**A**) For each sample, these plots show the distribution of genes detected per cell (nFeature_RNA), reads mapped per cell (nCount_RNA), and the percent of reads mapped to mitochondrial genes per cell (percent.mt), prior to any filtering. (**B**) Plots of the distributions of these QC parameters after filtering out cells with 1) fewer than 1000 genes detected, 2) greater than 7,000 genes detected, 3) greater than 50,000 mapped reads, and 4) greater than 5% mitochondrial reads. (C) Elbow plot of the standard deviations of each principal component (PC) of the t-cell population, based on the top 3,000 most variable genes. These values indicate how informative each PC is. This guided our choice of 30 PCs for subsequent clustering as PCs >30 contain little information. (**D**) Clustree plot of the T-cell population clustered at various resolutions from 0.5 to 5.0. Briefly, Clustree plots show how cells move between clusters as clustering resolution is increased, while the sc3 stability index indicates how stable a cluster is across all resolutions. This allows for rational selection of the resolution parameter. Each dot is a cluster. Each row corresponds to a resolution value, with values increasing from top to bottom. Dot size corresponds to the number of cells in the cluster. Arrows show how cells move from one cluster to another as resolution increases. Arrow color indicates the number of cells that move from cluster to cluster. Arrow transparency indicates the proportion of cells in a cluster that came from the source cluster at the previous resolution. Cluster color corresponds to sc3 stability, which indicates how stable a cluster is overall in tested resolutions. A final resolution of 1.0 (third row) was selected based on this plot, as clustering rapidly becomes unstable as the resolution increases much beyond this point.

**<u>Supplemental Figure S2:</u> Representative plots of the gating strategy used for flow cytometry experiments.** Multichromatic spectral flow cytometry was performed on uninfected and MRSA-infected BLT mice. **(A)** Gating strategy used on thawed/unstimulated cells to evaluate differences in CD4 T cells. Plots shown are using bone marrow cells. (B) Gating strategy used on stimulated cells to evaluate differences in cytokine production in activated cells. Plots shown are using bone marrow cells.

**<u>Supplemental Figure S3:</u> Increased number of activated cytokine producing CD4+ T cells in the bone marrow from MRSA-infected tibiae.** Post-stimulation with PMA/ionomycin live human CD45+/CD3+/CD4+/CD69+ T cells expressing checkpoint molecules were probed for functional capacity using the cytokines IFN-!, TNFɑ, IL-17A, and IL-2 (n=4-6 mice, *p<0.05, ANOVA).

**<u>Supplemental Figure S4:</u> Examination of bone marrow CD4 T cells expressing TIM-3 and LAG-3 for their functional capacity.** Post-stimulation with PMA/ionomycin live human CD45+/CD3+/CD4+/CD69+ T cells expressing checkpoint molecules were probed for functional capacity using the cytokines IFN-!, TNFɑ, IL-17A, and IL-2. Note that TIM-3+ and LAG-3+ CD4 T cells generally have diminished cytokine-secreting abilities, suggesting dysfunction (n=4-6 mice, *p<0.05, ANOVA).

**<u>Supplemental Figure S5:</u> Proliferation of immune cells is impaired in Hu-BLT mice infected with *S. aureus*.** Spleen sections from non-infected and infected mice were stained with antibodies specific for CD3 (red) and PCNA (white). Nuclei were labeled with DAPI. A) Spleen from sham-infected Hu-BLT mice show increased proliferating T cells. B) Spleen from Hu-BLT mice infected with *S. aureus* show a reduction in proliferating T cells. To estimate the proliferative activity in the spleens of Hu-BLT mice, the area covered by PCNA signal was measured with NIH Image J. C) Proliferation is significantly reduced in the spleens of *S. aureus* infected Hu-BLT mice, suggesting systemic immunosuppression (n=3, t-test, * p < 0.05).

**<u>Supplemental Figure S6:</u> Splenic CD4+ T cells expressing TIM-3 and LAG3 checkpoint proteins exhibit diminished proliferative capacity and altered cytokine production due to *S. aureus* infection.** Multichromatic spectral flow cytometry on uninfected and MRSA-infected BLT mice tibial bone marrow cells was performed **(A)** on unstimulated cells **(B)** post-stimulation with PMA/ionomycin. **(A)** Live human CD45+/CD3+/CD4+ T cells expressing checkpoint molecules TIM-3, LAG3, and PD-1 were probed for their proliferative capacity using the cell surface marker Ki67. Note that CD4+TIM-3+ and CD4+LAG3+ cells have lower amounts of proliferating Ki67+ cells in the bone marrow of infected BLT mice, suggesting functional exhaustion and dysfunction. **(B)** Live human CD45+/CD3+/CD4+/CD69+ T cells expressing checkpoint molecules TIM-3 and LAG-3 were probed for functional capacity using the cytokines IFN-!, TNFɑ, IL-17A, and IL-2 (n=4-9 mice, *p<0.05, ANOVA).

**Supplemental Table 1.**
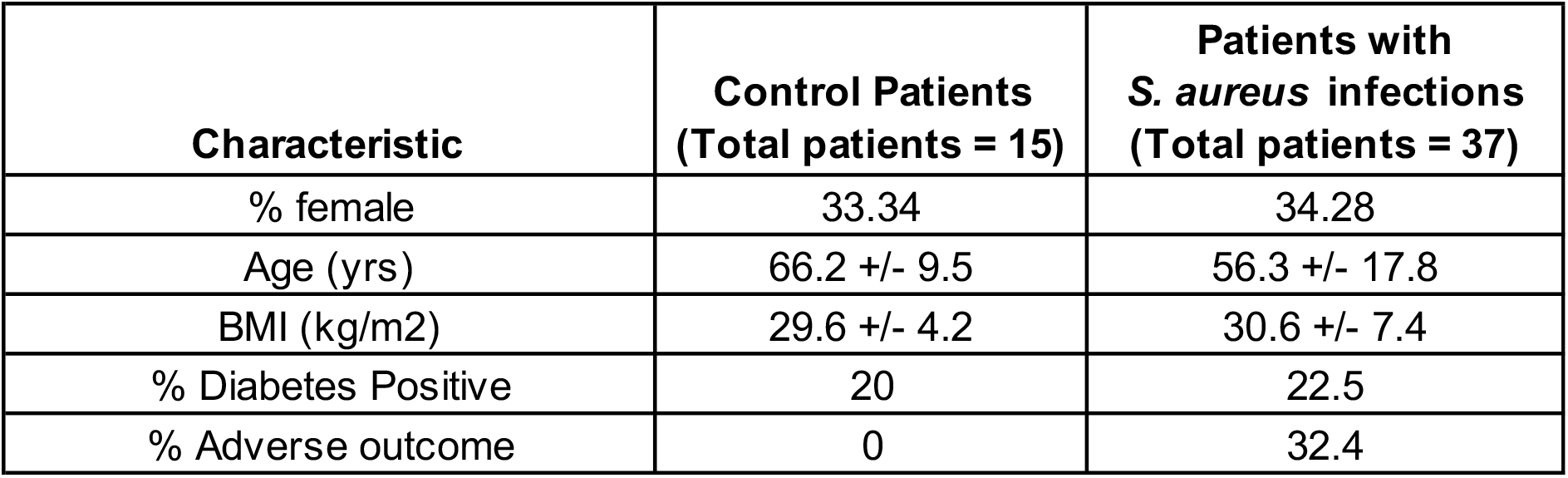
Demographic and outcome data of patients enrolled in the clinical study.

## References

1 Schwarz, E. M. et al. 2018 International Consensus Meeting on Musculoskeletal Infection: Research Priorities from the General Assembly Questions. J Orthop Res 37, 997–1006 (2019). 10.1002/jor.24293

2 Tande, A. J. & Patel, R. Prosthetic joint infection. Clin Microbiol Rev 27, 302–345 (2014). 10.1128/CMR.00111-13

3 Van Hal, S. J. et al. Predictors of Mortality in Staphylococcus aureus Bacteremia. Clinical Microbiology Reviews 25, 362–386 (2012). 10.1128/cmr.05022-11

4 Cram, P. et al. Total knee arthroplasty volume, utilization, and outcomes among Medicare beneficiaries, 1991-2010. Jama 308, 1227–1236 (2012). 10.1001/2012.jama.11153

5 Masters, E. A. et al. Evolving concepts in bone infection: redefining "biofilm", "acute vs. chronic osteomyelitis", "the immune proteome" and "local antibiotic therapy". Bone Res 7, 20 (2019). 10.1038/s41413-019-0061-z

6 Masters, E. A. et al. Skeletal infections: microbial pathogenesis, immunity and clinical management. Nature reviews. Microbiology 20, 385–400 (2022). 10.1038/s41579-022-00686-0

7 Libraty, D. H., Patkar, C. & Torres, B. *Staphylococcus aureus*Reactivation Osteomyelitis after 75 Years. New England Journal of Medicine 366, 481–482 (2012). 10.1056/nejmc1111493

8 Schwarz, E. M. et al. The 2023 Orthopaedic Research Society’s International Consensus Meeting on musculoskeletal infection: Summary from the host immunity section. J Orthop Res 42, 518–530 (2024). 10.1002/jor.25758

9 Kaech, S. M., Wherry, E. J. & Ahmed, R. Effector and memory T-cell differentiation: implications for vaccine development. Nature Reviews Immunology 2, 251–262 (2002). 10.1038/nri778

10 Caza, T. & Landas, S. Functional and Phenotypic Plasticity of CD4+ T Cell Subsets. BioMed Research International 2015, 1–13 (2015). 10.1155/2015/521957

11 Broker, B. M., Mrochen, D. & Peton, V. The T Cell Response to Staphylococcus aureus. Pathogens 5 (2016). 10.3390/pathogens5010031

12 Miggelbrink, A. M. et al. CD4 T-Cell Exhaustion: Does It Exist and What Are Its Roles in Cancer? Clin Cancer Res 27, 5742–5752 (2021). 10.1158/1078-0432.CCR-21-0206

13 Sokhi, U. K. et al. Immune Response to Persistent Staphyloccocus Aureus Periprosthetic Joint Infection in a Mouse Tibial Implant Model. J Bone Miner Res (2021). 10.1002/jbmr.4489

14 Muthukrishnan, G. et al. Humanized Mice Exhibit Exacerbated Abscess Formation and Osteolysis During the Establishment of Implant-Associated Staphylococcus aureus Osteomyelitis. Front Immunol 12 (2021). 10.3389/fimmu.2021.651515

15 Holtfreter, S. & Broker, B. M. Staphylococcal superantigens: do they play a role in sepsis? Arch Immunol Ther Exp (Warsz) 53, 13–27 (2005).

16 Grumann, D. et al. Immune cell activation by enterotoxin gene cluster (egc)-encoded and non-egc superantigens from Staphylococcus aureus. J Immunol 181, 5054–5061 (2008). 10.4049/jimmunol.181.7.5054

17 Grumann, D., Nubel, U. & Broker, B. M. Staphylococcus aureus toxins--their functions and genetics. Infect Genet Evol 21, 583–592 (2014). 10.1016/j.meegid.2013.03.013

18 Oliveira, D., Borges, A. & Simoes, M. Staphylococcus aureus Toxins and Their Molecular Activity in Infectious Diseases. Toxins (Basel) 10 (2018). 10.3390/toxins10060252

19 Wherry, E. J. & Kurachi, M. Molecular and cellular insights into T cell exhaustion. Nature reviews. Immunology 15, 486–499 (2015). 10.1038/nri3862

20 Blank, C. U. et al. Defining ‘T cell exhaustion’. Nature reviews. Immunology 19, 665–674 (2019). 10.1038/s41577-019-0221-9

21 Saeidi, A. et al. T-Cell Exhaustion in Chronic Infections: Reversing the State of Exhaustion and Reinvigorating Optimal Protective Immune Responses. Front Immunol 9, 2569 (2018). 10.3389/fimmu.2018.02569

22 Baessler, A. & Vignali, D. A. A. T Cell Exhaustion. Annu Rev Immunol (2024). 10.1146/annurev-immunol-090222-110914

23 Silberstein, J. L. et al. Structural insights reveal interplay between LAG-3 homodimerization, ligand binding, and function. Proceedings of the National Academy of Sciences 121 (2024). 10.1073/pnas.2310866121

24 Keane, C. et al. LAG3: a novel immune checkpoint expressed by multiple lymphocyte subsets in diffuse large B-cell lymphoma. Blood Advances 4, 1367–1377 (2020). 10.1182/bloodadvances.2019001390

25 Jin, H.-T. et al. Cooperation of Tim-3 and PD-1 in CD8 T-cell exhaustion during chronic viral infection. Proceedings of the National Academy of Sciences 107, 14733–14738 (2010). 10.1073/pnas.1009731107

26 Jones, R. B. et al. Tim-3 expression defines a novel population of dysfunctional T cells with highly elevated frequencies in progressive HIV-1 infection. The Journal of Experimental Medicine 205, 2763–2779 (2008). 10.1084/jem.20081398

27 Day, C. L. et al. PD-1 expression on HIV-specific T cells is associated with T-cell exhaustion and disease progression. Nature 443, 350–354 (2006). 10.1038/nature05115

28 Barber, D. L. et al. Restoring function in exhausted CD8 T cells during chronic viral infection. Nature 439, 682–687 (2006). 10.1038/nature04444

29 Elahi, S., Shahbaz, S. & Houston, S. Selective Upregulation of CTLA-4 on CD8+ T Cells Restricted by HLA-B*35Px Renders them to an Exhausted Phenotype in HIV-1 infection. PLOS Pathogens 16, e1008696 (2020). 10.1371/journal.ppat.1008696

30 Kaufmann, D. E. et al. Upregulation of CTLA-4 by HIV-specific CD4+ T cells correlates with disease progression and defines a reversible immune dysfunction. Nature Immunology 8, 1246–1254 (2007). 10.1038/ni1515

31 Li, H., Llera, A., Malchiodi, E. L. & Mariuzza, R. A. The structural basis of T cell activation by superantigens. Annu Rev Immunol 17, 435–466 (1999). 10.1146/annurev.immunol.17.1.435

32 Proft, T. & Fraser, J. D. Bacterial superantigens. Clin Exp Immunol 133, 299–306 (2003). 10.1046/j.1365-2249.2003.02203.x

33 Spaulding, A. R. et al. Staphylococcal and streptococcal superantigen exotoxins. Clinical microbiology reviews 26, 422–447 (2013). 10.1128/CMR.00104-12

34 Kim, H. K., Thammavongsa, V., Schneewind, O. & Missiakas, D. Recurrent infections and immune evasion strategies of Staphylococcus aureus. Curr Opin Microbiol 15, 92–99 (2012). 10.1016/j.mib.2011.10.012

35 Watson, A. R., Janik, D. K. & Lee, W. T. Superantigen-induced CD4 memory T cell anergy. I. Staphylococcal enterotoxin B induces Fyn-mediated negative signaling. Cell Immunol 276, 16–25 (2012). 10.1016/j.cellimm.2012.02.003

36 Proctor, R. A. Immunity to Staphylococcus aureus: Implications for Vaccine Development. Microbiol Spectr 7 (2019). 10.1128/microbiolspec.GPP3-0037-2018

37 Janik, D. K. & Lee, W. T. Staphylococcal Enterotoxin B (SEB) Induces Memory CD4 T Cell Anergy in vivo and Impairs Recall Immunity to Unrelated Antigens. J Clin Cell Immunol 6, 1–8 (2015). 10.4172/2155-9899.1000346

38 Chen, J. et al. The development and improvement of immunodeficient mice and humanized immune system mouse models. Front Immunol 13, 1007579 (2022). 10.3389/fimmu.2022.1007579

39 Shultz, L. D., Brehm, M. A., Garcia-Martinez, J. V. & Greiner, D. L. Humanized mice for immune system investigation: progress, promise and challenges. Nat Rev Immunol 12, 786–798 (2012). 10.1038/nri3311

40 Lee, J. Y., Han, A. R. & Lee, D. R. T Lymphocyte Development and Activation in Humanized Mouse Model. Dev Reprod 23, 79–92 (2019). 10.12717/DR.2019.23.2.079

41 Billerbeck, E. et al. Development of human CD4+FoxP3+ regulatory T cells in human stem cell factor-, granulocyte-macrophage colony-stimulating factor-, and interleukin-3-expressing NOD-SCID IL2Rgamma(null) humanized mice. Blood 117, 3076–3086 (2011). 10.1182/blood-2010-08-301507

42 Wunderlich, M. et al. AML xenograft efficiency is significantly improved in NOD/SCID-IL2RG mice constitutively expressing human SCF, GM-CSF and IL-3. Leukemia 24, 1785–1788 (2010). 10.1038/leu.2010.158

43 Li, D. et al. Quantitative mouse model of implant-associated osteomyelitis and the kinetics of microbial growth, osteolysis, and humoral immunity. J Orthop Res 26, 96–105 (2008). 10.1002/jor.20452

44 Varrone, J. J. et al. Passive immunization with anti-glucosaminidase monoclonal antibodies protects mice from implant-associated osteomyelitis by mediating opsonophagocytosis of Staphylococcus aureus megaclusters. J Orthop Res 32, 1389–1396 (2014). 10.1002/jor.22672

45 Nishitani, K. et al. Quantifying the natural history of biofilm formation in vivo during the establishment of chronic implant-associated Staphylococcus aureus osteomyelitis in mice to identify critical pathogen and host factors. J Orthop Res 33, 1311–1319 (2015). 10.1002/jor.22907

46 De Mesy Bentley, K. L., et al. Evidence of Staphylococcus Aureus Deformation, Proliferation, and Migration in Canaliculi of Live Cortical Bone in Murine Models of Osteomyelitis. J Bone Miner Res 32, 985–990 (2017). 10.1002/jbmr.3055

47 Morita, Y. et al. Systemic IL-27 administration prevents abscess formation and osteolysis via local neutrophil recruitment and activation. Bone Research 10, 56 (2022). 10.1038/s41413-022-00228-7

48 Butler, A., Hoffman, P., Smibert, P., Papalexi, E. & Satija, R. Integrating single-cell transcriptomic data across different conditions, technologies, and species. Nat Biotechnol 36, 411–420 (2018). 10.1038/nbt.4096

49 Hao, Y. et al. Integrated analysis of multimodal single-cell data. Cell 184, 3573–3587 e3529 (2021). 10.1016/j.cell.2021.04.048

50 Satija, R., Farrell, J. A., Gennert, D., Schier, A. F. & Regev, A. Spatial reconstruction of single-cell gene expression data. Nat Biotechnol 33, 495–502 (2015). 10.1038/nbt.3192

51 Stuart, T. et al. Comprehensive Integration of Single-Cell Data. Cell 177, 1888–1902 e1821 (2019). 10.1016/j.cell.2019.05.031

52 Becht, E. et al. Dimensionality reduction for visualizing single-cell data using UMAP. Nat Biotechnol (2018). 10.1038/nbt.4314

53 Im, S. J. et al. Defining CD8+ T cells that provide the proliferative burst after PD-1 therapy. Nature 537, 417–421 (2016). 10.1038/nature19330

54 Muthukrishnan, G. et al. Serum antibodies against Staphylococcus aureus can prognose treatment success in patients with bone infections. J Orthop Res 39, 2169–2176 (2021). 10.1002/jor.24955

55 Cao, Y. et al. Risk stratification biomarkers for Staphylococcus aureus bacteraemia. Clin Transl Immunology 9, e1110 (2020). 10.1002/cti2.1110

56 Wozniak, J. M. et al. Mortality Risk Profiling of Staphylococcus aureus Bacteremia by Multi-omic Serum Analysis Reveals Early Predictive and Pathogenic Signatures. Cell 182, 1311–1327.e1314 (2020). 10.1016/j.cell.2020.07.040

57 Knop, J. et al. Staphylococcus aureus Infection in Humanized Mice: A New Model to Study Pathogenicity Associated With Human Immune Response. The Journal of infectious diseases 212, 435–444 (2015). 10.1093/infdis/jiv073

58 Tseng, C. W. et al. Increased Susceptibility of Humanized NSG Mice to Panton-Valentine Leukocidin and Staphylococcus aureus Skin Infection. PLoS pathogens 11, e1005292 (2015). 10.1371/journal.ppat.1005292

59 Prince, A., Wang, H., Kitur, K. & Parker, D. Humanized Mice Exhibit Increased Susceptibility to Staphylococcus aureus Pneumonia. The Journal of infectious diseases 215, 1386–1395 (2017). 10.1093/infdis/jiw425

60 Hung, S. et al. Next-generation humanized NSG-SGM3 mice are highly susceptible to Staphylococcus aureus infection. Front Immunol 14, 1127709 (2023). 10.3389/fimmu.2023.1127709

61 Hofstee, M. I. et al. Staphylococcus aureus Panton-Valentine Leukocidin worsens acute implant-associated osteomyelitis in humanized BRGSF mice. JBMR Plus (2024). 10.1093/jbmrpl/ziad005

62 Alonzo, F., 3rd & Torres, V. J. Bacterial survival amidst an immune onslaught: the contribution of the Staphylococcus aureus leukotoxins. PLoS Pathog 9, e1003143 (2013). 10.1371/journal.ppat.1003143

63 Alonzo, F., 3rd & Torres, V. J. The bicomponent pore-forming leucocidins of Staphylococcus aureus. Microbiol Mol Biol Rev 78, 199–230 (2014). 10.1128/MMBR.00055-13

64 Petersson, K., Pettersson, H., Skartved, N. J., Walse, B. & Forsberg, G. Staphylococcal enterotoxin H induces V alpha-specific expansion of T cells. J Immunol 170, 4148–4154 (2003). 10.4049/jimmunol.170.8.4148

65 Li, S. J. et al. Superantigenic activity of toxic shock syndrome toxin-1 is resistant to heating and digestive enzymes. J Appl Microbiol 110, 729–736 (2011). 10.1111/j.1365-2672.2010.04927.x

66 Tebartz, C. et al. A Major Role for Myeloid-Derived Suppressor Cells and a Minor Role for Regulatory T Cells in Immunosuppression during *Staphylococcus aureus* Infection. The Journal of Immunology 194, 1100–1111 (2015). 10.4049/jimmunol.1400196

67 Zhu, L. et al. The correlation between the Th17/Treg cell balance and bone health. Immunity & Ageing 17 (2020). 10.1186/s12979-020-00202-z

68 Pan, J. et al. Landscape of Exhausted T Cells in Tuberculosis Revealed by Single-Cell Sequencing. Microbiology Spectrum 11 (2023). 10.1128/spectrum.02839-22

69 Van Roy, Z. et al. Tissue niche influences immune and metabolic profiles to Staphylococcus aureus biofilm infection. Nat Commun 15, 8965 (2024). 10.1038/s41467-024-53353-8

70 Surendar, J. et al. Osteomyelitis is associated with increased anti-inflammatory response and immune exhaustion. Front Immunol 15, 1396592 (2024). 10.3389/fimmu.2024.1396592

71 Wilde, A. D. et al. Bacterial Hypoxic Responses Revealed as Critical Determinants of the Host-Pathogen Outcome by TnSeq Analysis of Staphylococcus aureus Invasive Infection. PLOS Pathogens 11, e1005341 (2015). 10.1371/journal.ppat.1005341

72 Heim, C. E. et al. Human prosthetic joint infections are associated with myeloid-derived suppressor cells (MDSCs): Implications for infection persistence. J Orthop Res 36, 1605–1613 (2018). 10.1002/jor.23806

73 Heim, C. E. et al. Myeloid-derived suppressor cells contribute to Staphylococcus aureus orthopedic biofilm infection. J Immunol 192, 3778–3792 (2014). 10.4049/jimmunol.1303408

74 van der Windt, G. J. & Pearce, E. L. Metabolic switching and fuel choice during T-cell differentiation and memory development. Immunol Rev 249, 27–42 (2012). 10.1111/j.1600-065X.2012.01150.x

75 Geltink, R. I. K., Kyle, R. L. & Pearce, E. L. Unraveling the Complex Interplay Between T Cell Metabolism and Function. Annu Rev Immunol 36, 461–488 (2018). 10.1146/annurev-immunol-042617-053019

76 Kersten, K. et al. Spatiotemporal co-dependency between macrophages and exhausted CD8(+) T cells in cancer. Cancer Cell 40, 624–638 e629 (2022). 10.1016/j.ccell.2022.05.004

77 Zhang, W. et al. Staphylococcus aureus infection initiates hypoxia mediated transforming growth factor β1 upregulation to trigger osteomyelitis. mSystems 7 (2022). 10.1128/msystems.00380-22

78 Wang, Y. T. et al. Metabolic adaptation supports enhanced macrophage efferocytosis in limited-oxygen environments. Cell Metab 35, 316–331 e316 (2023). 10.1016/j.cmet.2022.12.005

79 Yin, C. & Heit, B. Cellular Responses to the Efferocytosis of Apoptotic Cells. Front Immunol 12, 631714 (2021). 10.3389/fimmu.2021.631714

80 Lowther, D. E. et al. PD-1 marks dysfunctional regulatory T cells in malignant gliomas. JCI Insight 1 (2016). 10.1172/jci.insight.85935

81 McLane, L. M., Abdel-Hakeem, M. S. & Wherry, E. J. CD8 T Cell Exhaustion During Chronic Viral Infection and Cancer. Annual Review of Immunology 37, 457–495 (2019). 10.1146/annurev-immunol-041015-055318

82 Han, S., Asoyan, A., Rabenstein, H., Nakano, N. & Obst, R. Role of antigen persistence and dose for CD4+ T-cell exhaustion and recovery. Proceedings of the National Academy of Sciences of the United States of America 107, 20453–20458 (2010). 10.1073/pnas.1008437107

83 Wildeman, P. et al. Genomic characterization and outcome of prosthetic joint infections caused by Staphylococcus aureus. Sci Rep 10, 5938 (2020). 10.1038/s41598-020-62751-z

84 Sun, Q. et al. Immune checkpoint therapy for solid tumours: clinical dilemmas and future trends. Signal Transduction and Targeted Therapy 8 (2023). 10.1038/s41392-023-01522-4

85 Li, K. et al. PD-1/PD-L1 blockade is a potent adjuvant in treatment of Staphylococcus aureus osteomyelitis in mice. Mol Ther 31, 174–192 (2023). 10.1016/j.ymthe.2022.09.006

86 Biedermann, L. et al. Inflammation of Bone in Patients with Periprosthetic Joint Infections of the Knee. JB JS Open Access 8 (2023). 10.2106/JBJS.OA.22.00101

87 Parlet, C. P., Brown, M. M. & Horswill, A. R. Commensal Staphylococci Influence Staphylococcus aureus Skin Colonization and Disease. Trends Microbiol 27, 497–507 (2019). 10.1016/j.tim.2019.01.008

88 Hendriks, A. et al. Staphylococcus aureus-Specific Tissue-Resident Memory CD4(+) T Cells Are Abundant in Healthy Human Skin. Front Immunol 12, 642711 (2021). 10.3389/fimmu.2021.642711

89 Ando, M., Ito, M., Srirat, T., Kondo, T. & Yoshimura, A. Memory T cell, exhaustion, and tumor immunity. Immunol Med 43, 1–9 (2020). 10.1080/25785826.2019.1698261

90 Morgenstern, M. et al. The AO trauma CPP bone infection registry: Epidemiology and outcomes of Staphylococcus aureus bone infection. J Orthop Res (2020). 10.1002/jor.24804

91 Campbell, M. P. et al. Low albumin level is more strongly associated with adverse outcomes and Staphylococcus aureus infection than hemoglobin A1C or smoking tobacco. J Orthop Res (2022). 10.1002/jor.25282

92 Lan, P. et al. Induction of human T-cell tolerance to porcine xenoantigens through mixed hematopoietic chimerism. Blood 103, 3964–3969 (2004). 10.1182/blood-2003-10-3697

93 Onoe, T. et al. Human natural regulatory T cell development, suppressive function, and postthymic maturation in a humanized mouse model. J Immunol 187, 3895–3903 (2011). 10.4049/jimmunol.1100394

94 Kalscheuer, H. et al. A model for personalized in vivo analysis of human immune responsiveness. Sci Transl Med 4, 125ra130 (2012). 10.1126/scitranslmed.3003481

95 Botros, M. et al. Cutibacterium acnes invades submicron osteocyte lacuno-canalicular networks following implant-associated osteomyelitis. J Orthop Res 42, 2593–2603 (2024). 10.1002/jor.25929

96 Nishitani, K. et al. IsdB antibody-mediated sepsis following S. aureus surgical site infection. JCI Insight 5 (2020). 10.1172/jci.insight.141164

97 Masters, E. A. et al. Identification of Penicillin Binding Protein 4 (PBP4) as a critical factor for Staphylococcus aureus bone invasion during osteomyelitis in mice. PLoS pathogens 16, e1008988 (2020). 10.1371/journal.ppat.1008988

98 Zappia, L. & Oshlack, A. Clustering trees: a visualization for evaluating clusterings at multiple resolutions. Gigascience 7 (2018). 10.1093/gigascience/giy083

99 Aran, D. et al. Reference-based analysis of lung single-cell sequencing reveals a transitional profibrotic macrophage. Nat Immunol 20, 163–172 (2019). 10.1038/s41590-018-0276-y

100 Hafemeister, C. & Satija, R. Normalization and variance stabilization of single-cell RNA-seq data using regularized negative binomial regression. Genome Biol 20, 296 (2019). 10.1186/s13059-019-1874-1

