## Supplemental Figures for "Immune Checkpoint Molecules as Biomarkers of *Staphylococcus aureus* Bone Infection and Clinical Outcome"

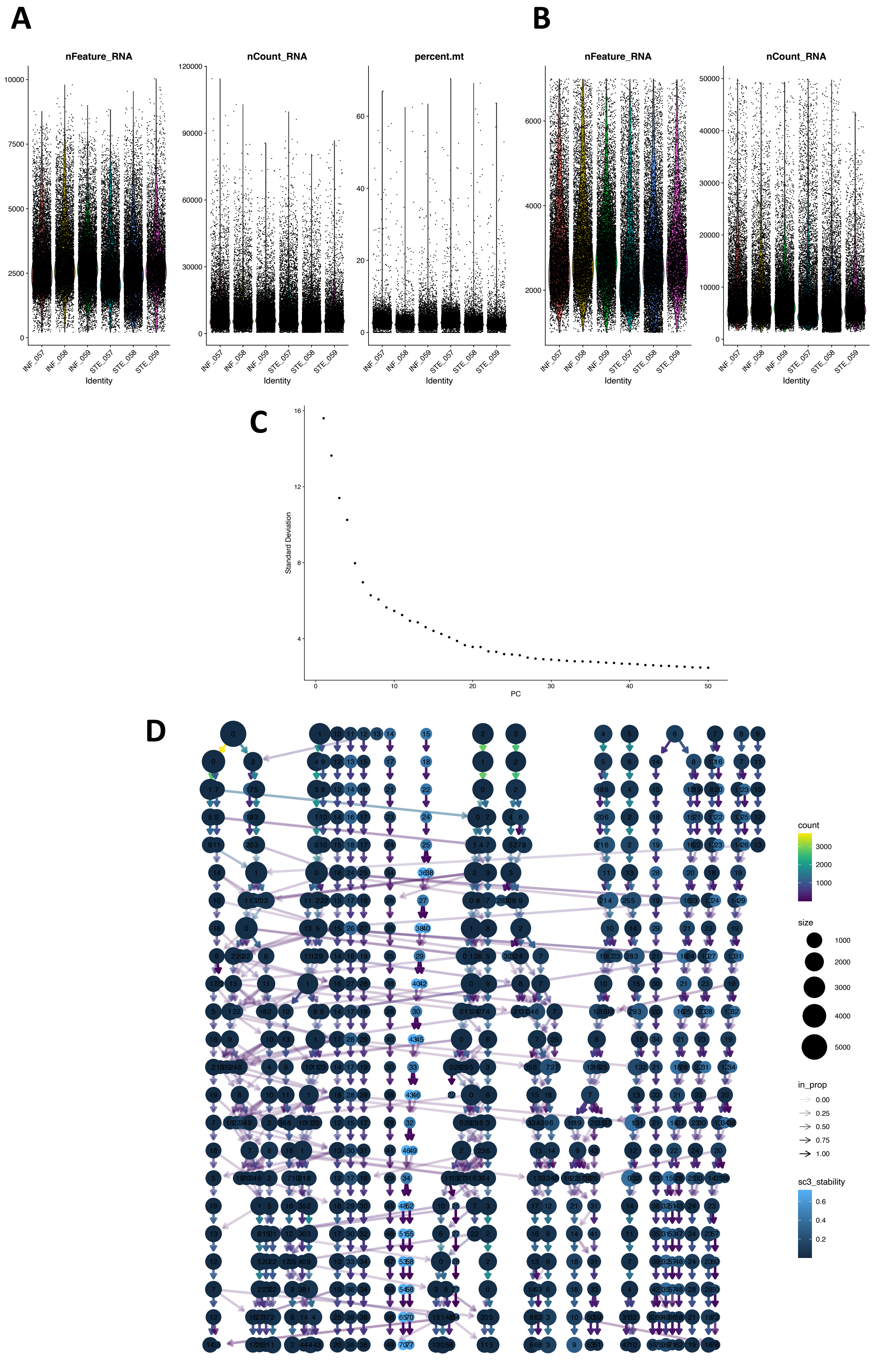

A
B
nFeature_RNA
nCount_RNA
percent.mt
INF_057
INF_058
INF_059
STE_057
STE_058
STE_059
Identity
C
Standard Deviation
PC
D
count
size
in_prop
sc3_stability

**<u>Supplementary Figure S1.</u> Quality Control, Dimension Reduction, and Clustering Overview.** (**A**) For each sample, these plots show the distribution of genes detected per cell (nFeature_RNA), reads mapped per cell (nCount_RNA), and the percent of reads mapped to mitochondrial genes per cell (percent.mt), prior to any filtering. (**B**) Plots of the distributions of these QC parameters after filtering out cells with 1) fewer than 1000 genes detected, 2) greater than 7,000 genes detected, 3) greater than 50,000 mapped reads, and 4) greater than 5% mitochondrial reads. (C**)** Elbow plot of the standard deviations of each principal component (PC) of the t-cell population, based on the top 3,000 most variable genes. These values indicate how informative each PC is. This guided our choice of 30 PCs for subsequent clustering as PCs >30 contain little information. (**D**) Clustree plot of the T-cell population clustered at various resolutions from 0.5 to 5.0. Briefly, Clustree plots show how cells move between clusters as clustering resolution is increased, while the sc3 stability index indicates how stable a cluster is across all resolutions. This allows for rational selection of the resolution parameter. Each dot is a cluster. Each row corresponds to a resolution value, with values increasing from top to bottom. Dot size corresponds to the number of cells in the cluster. Arrows show how cells move from one cluster to another as resolution increases. Arrow color indicates the number of cells that move from cluster to cluster. Arrow transparency indicates the proportion of cells in a cluster that came from the source cluster at the previous resolution. Cluster color corresponds to sc3 stability, which indicates how stable a cluster is over all tested resolutions. A final resolution of 1.0 (third row) was selected based on this plot, as clustering rapidly becomes unstable as the resolution increases much beyond this point.

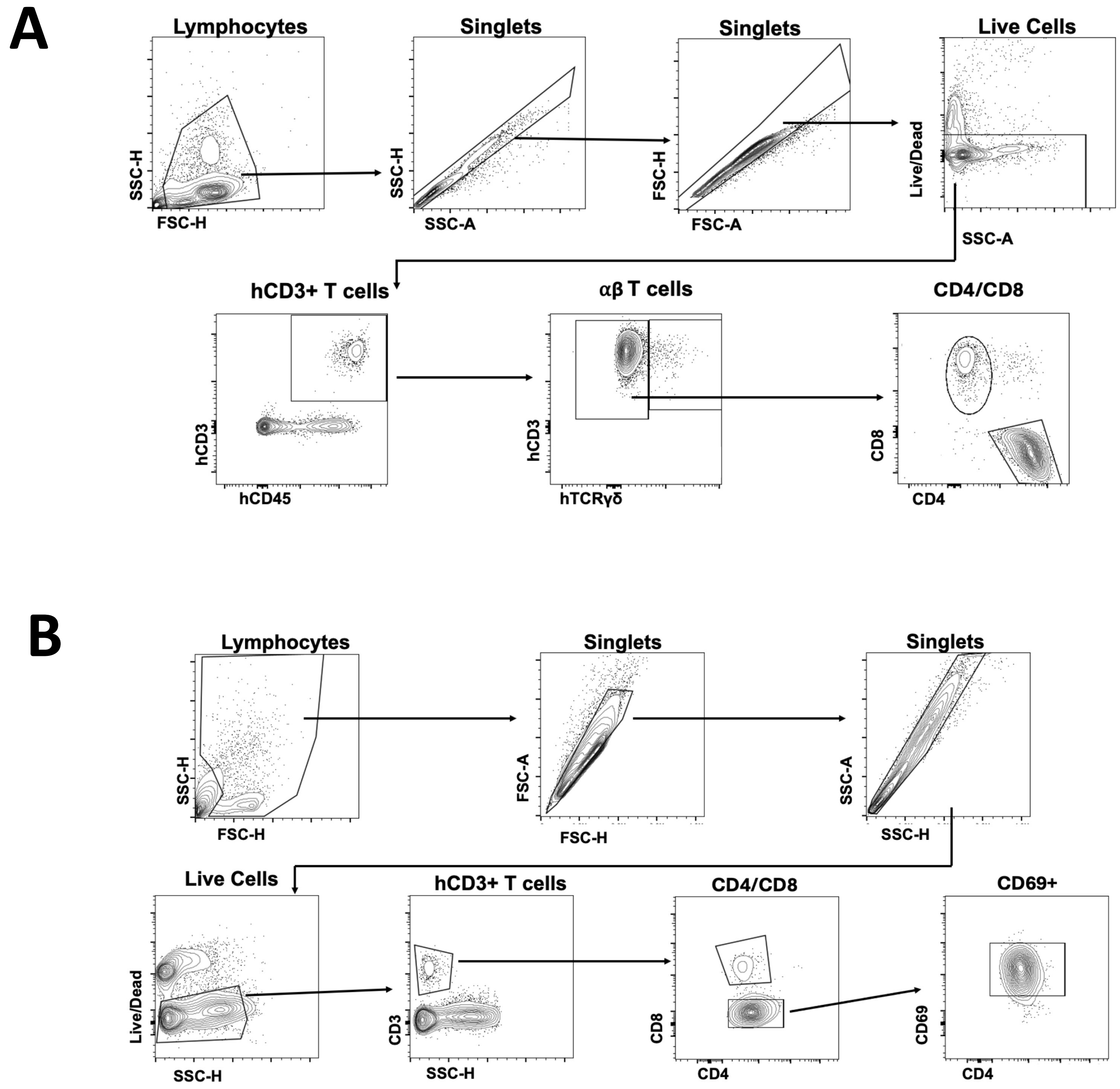


**Supplemental Figure S2: Representative plots of the gating strategy used for flow cytometry experiments.** Multichromatic spectral flow cytometry was performed on uninfected and MRSA-infected BLT mice. **(A)** Gating strategy used on thawed/unstimulated cells to evaluate differences in CD4 T cells. Plots shown are using bone marrow cells. (B) Gating strategy used on stimulated cells to evaluate differences in cytokine production in activated cells. Plots shown are using bone marrow cells.

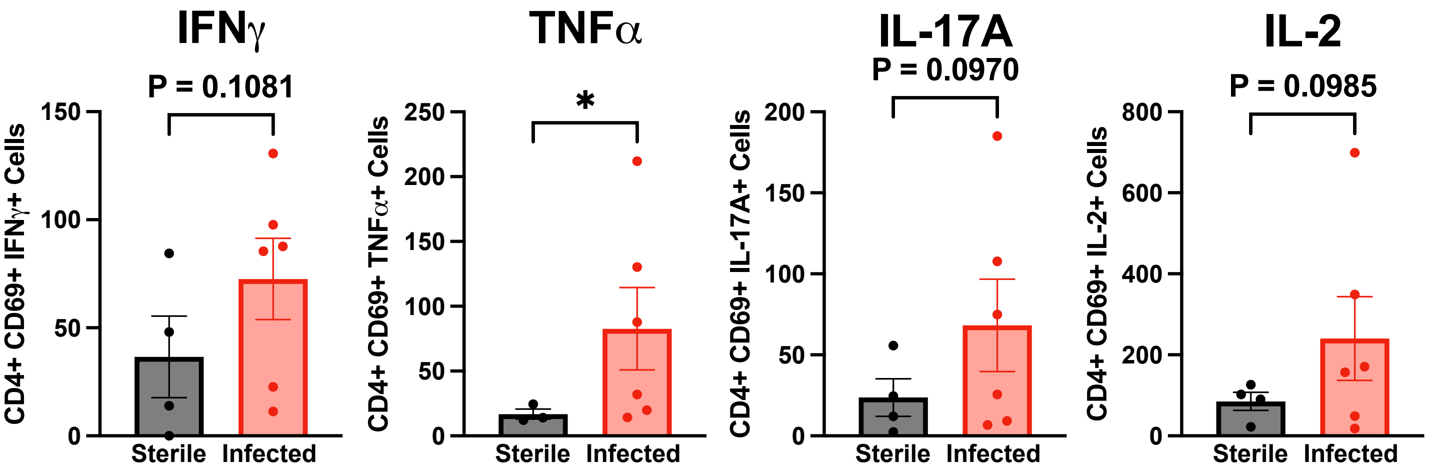


**Supplemental Figure S3: Increased number of activated cytokine producing CD4+ T cells in the bone marrow from MRSA-infected tibiae.** Post-stimulation with PMA/ionomycin live human CD45+/CD3+/CD4+/CD69+ T cells expressing checkpoint molecules were probed for functional capacity using the cytokines IFN-**γ**, TNFα, IL-17A, and IL-2 (n=4-6 mice, $*p<0.05$, ANOVA).

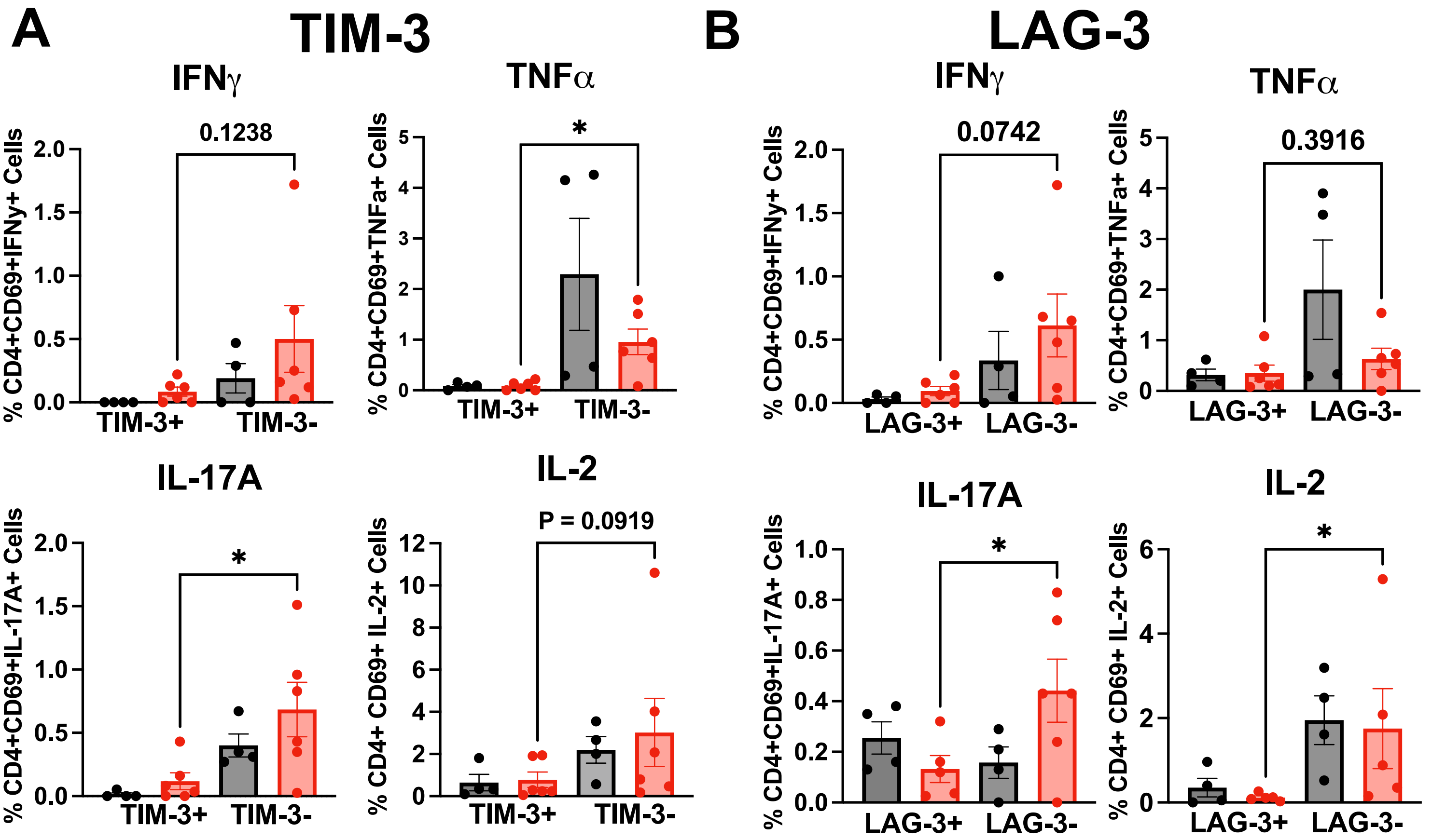


**<u>Supplemental Figure S4:</u> Examination of bone marrow CD4 T cells expressing TIM-3 and LAG-3 for their functional capacity**. Post-stimulation with PMA/ionomycin live human CD45+/CD3+/CD4+/CD69+ T cells expressing checkpoint molecules were probed for functional capacity using the cytokines IFN-**γ**, TNFα, IL-17A, and IL-2. Note that TIM-3+ and LAG-3+ CD4 T cells generally have diminished cytokine-secreting abilities, suggesting dysfunction (n=4-6 mice, *p<0.05, ANOVA).

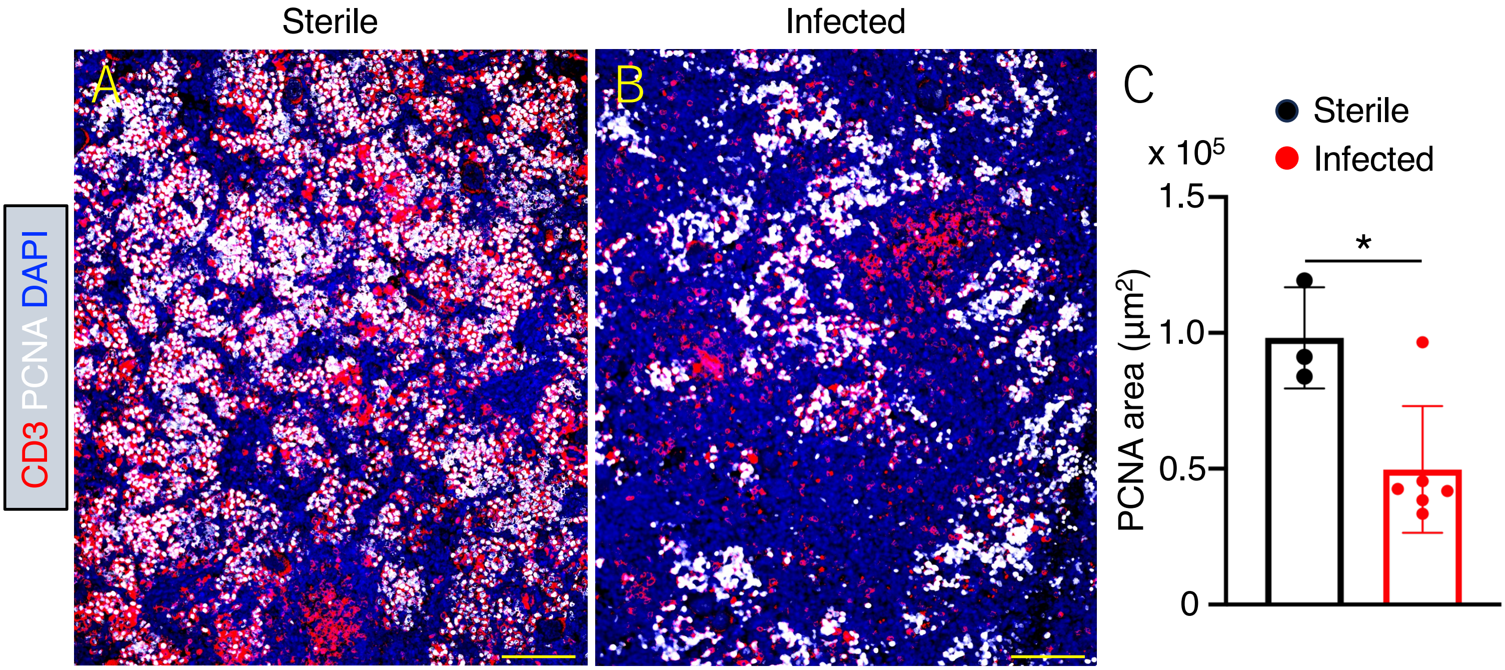


**Supplemental Figure S5. Proliferation of immune cells is impaired in Hu-BLT mice infected with *S. aureus*.** Spleen sections from non-infected and infected mice were stained with antibodies specific for CD3 (red) and PCNA (white). Nuclei were labeled with DAPI. (**A**) Spleen from sham-infected Hu-BLT mice show increased proliferating T cells. (**B**) Spleen from Hu-BLT mice infected with *S. aureus* show a reduction in proliferating T cells. To estimate the proliferative activity in the spleens of Hu-BLT mice, the area covered by PCNA signal was measured with NIH Image J. (**C**) Proliferation is significantly reduced in the spleens of *S. aureus* infected Hu-BLT mice suggesting systemic immunosuppression (n=3, t-test, * $p < 0.05$).

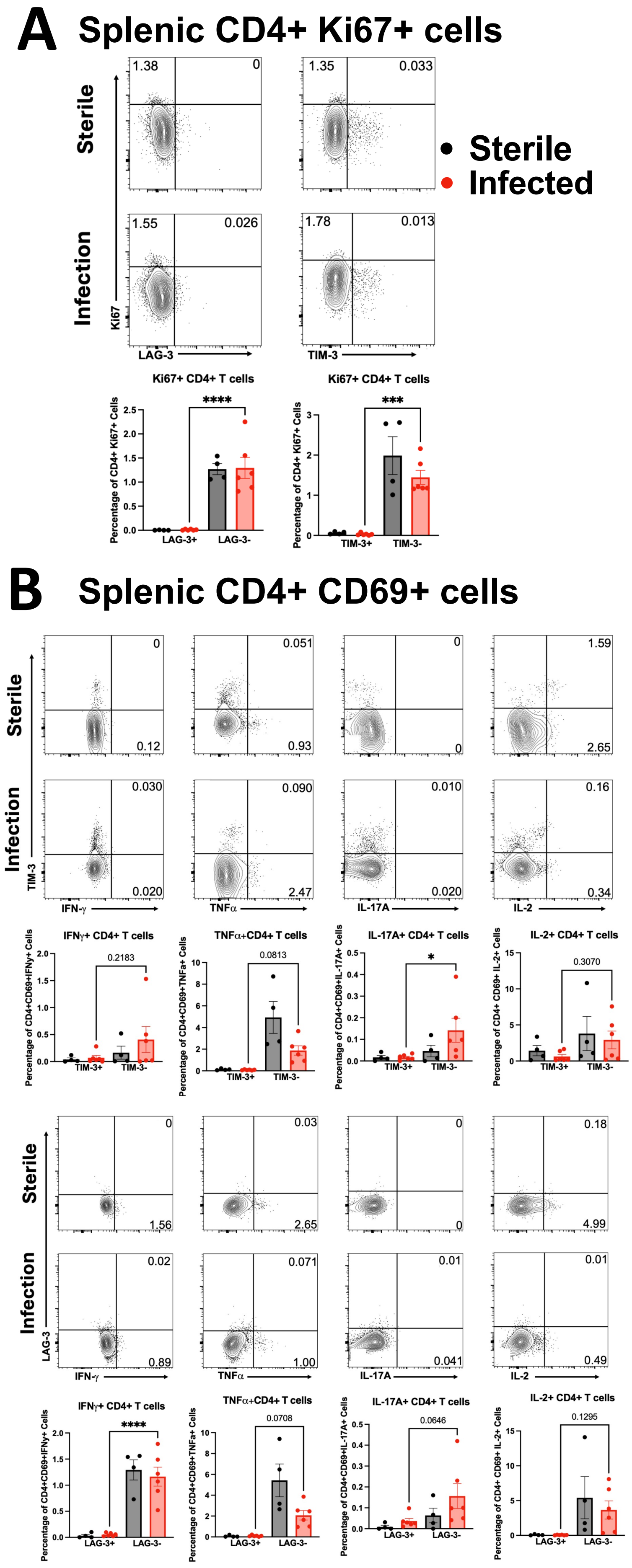


**<u>Supplemental Figure S6:</u> Splenic CD4+ T cells expressing TIM-3 and LAG3 checkpoint proteins exhibit diminished proliferative capacity and altered cytokine production due to *S. aureus* infection.** Multichromatic spectral flow cytometry on uninfected and MRSA-infected BLT mice tibial bone marrow cells was performed **(A)** on unstimulated cells **(B)** and stimulated cells post-stimulation with PMA/ionomycin. **(A)** Live human CD45+/CD3+/CD4+ T cells expressing checkpoint molecules TIM-3, LAG3, and PD-1 were probed for their proliferative capacity using the cell surface marker Ki67. Note that CD4+TIM-3+ and CD4+LAG3+ cells have lower amounts of proliferating Ki67+ cells in the bone marrow of infected BLT mice, suggesting functional exhaustion and dysfunction.

**(B)** Live human CD45+/CD3+/CD4+/CD69+ T cells expressing checkpoint molecules TIM-3 and LAG-3 were probed for functional capacity using the cytokines IFN-**γ**, TNFα, IL-17A, and IL-2 (n=4-9 mice, *$p<0.05$, ANOVA).
